# Deamidation disrupts native and transient contacts to weaken the interaction between UBC13 and RING-finger E3 ligases

**DOI:** 10.1101/665992

**Authors:** Priyesh Mohanty, Rashmi, Batul Ismail Habibullah, Arun G S, Ranabir Das

## Abstract

The deamidase OspI from enteric bacteria *Shigella flexneri* deamidates a glutamine residue in the host ubiquitin-conjugating enzyme UBC13 and converts it to glutamate (Q100E). Consequently, its polyubiquitination activity in complex with the RING-finger ubiquitin ligase TRAF6 and the downstream NF-κB inflammatory response is inactivated. The precise role of deamidation in inactivating the UBC13/TRAF6 complex is unknown. We report that deamidation inhibits the interaction between UBC13 and TRAF6 RING-domain (TRAF6^RING^) by perturbing both the native and transient interactions. Deamidation creates a new intramolecular salt-bridge in UBC13 that competes with a critical intermolecular salt-bridge at the native UBC13/TRAF6^RING^ interface. Moreover, the salt-bridge competition prevents transient interactions necessary to form a typical UBC13/RING complex. Repulsion between E100 and the negatively charged surface of RING also prevents transient interactions in the UBC13/RING complex. Our findings highlight a mechanism where a post-translational modification perturbs the conformation and stability of transient complexes to inhibit protein-protein association.

## Introduction

Several bacterial pathogens secrete effector proteins that inhibit or co-opt the Ubiquitin (Ub) pathway to suppress the immune response of the host cell (Ashida, Kim & Sasakawa 2014). The human pathogenic bacteria *Shigella flexneri* inactivates the host inflammatory Nf-κB signaling, responsible for inducing inflammatory cytokine responses during pathogen invasion (Sanada et al. 2012). In the early events of interleukin-dependent activation of Nf-κB signaling, the Ub-conjugating enzyme UBC13, and the Ub-ligase TRAF6 function together to synthesize both unanchored polyubiquitin chains, and anchored polyubiquitin chains on TRAF6 and its substrate NEMO (Chen 2005). These chains serve as a scaffold to bring together TAK1/2 and IKK kinases, eventually leading to phosphorylation and activation of the IKK kinases, IκB degradation and nuclear translocation of transcription factor Nf-κB. To inactivate Nf-κB signaling, *Shigella flexneri* secretes a Type III effector called OspI, which functions as a deamidase (Sanada et al. 2012). OspI specifically targets a glutamine residue in UBC13 and converts it to glutamate (Q100E). Ubiquitination reactions of TRAF6 either with UBC13 in the presence of OspI or with mutant Q100E-UBC13 (dUBC13) show a significant drop in polyubiquitination activity (Sanada et al. 2012). However, the mechanism underlying inhibition of polyubiquitination by deamidation of UBC13 remains unclear.

Ubiquitination is a eukaryotic post-translational modification (Komander & Rape 2012), wherein the last glycine residue in the C-terminal tail of Ubiquitin (Ub) is activated and covalently attached to a substrate lysine residue. Ubiquitination involves three steps: an initial activation and thioester conjugation by the Ubiquitin-activating enzyme (E1), followed by the thioester conjugation to the Ubiquitin-conjugating (E2) enzymes, and a final step in which the Ub is covalently attached to the substrate amino group. The last step is typically catalyzed by a class of Ubiquitin ligases (E3), which contain either a RING(Really Interesting New Gene)-finger domain, U-box domain or HECT domain (Metzger et al. 2010). The RING-finger/U-box domain stabilizes a catalytic-closed conformation of the flexible E2~Ub species and drastically enhances the rate of Ub conjugation to substrates (Dou et al. 2012; Plechanovov et al. 2012; Pruneda et al. 2012). The E2 UBC13 functions with several RING-finger E3s like TRAF6 to synthesize K63-linked poly-Ub chains that function to activate DNA repair or immune response (Matsuzawa et al. 2007). Apart from the E3s, UBC13 also binds a co-factor MMS2, which does not activate UBC13 but maintains the linkage specificity of the poly-Ub chains synthesized by UBC13 (Branigan et al. 2015).

In this study, we have investigated the mechanisms underlying the inactivation of UBC13 upon deamidation using NMR spectroscopy, molecular dynamics (MD) simulations, and *in-vitro* ubiquitination assays. We report that deamidation weakens the non-covalent interaction of UBC13 with RING-finger domain of TRAF6 (TRAF6^RING^), without perturbing UBC13 structure or the enzymatic activity of UBC13. However, the underlying cause of reduced interaction is nonintuitive since Q100 is in the vicinity of UBC13/TRAF6^RING^ interface but does not form any contact with the TRAF6^RING^. Further studies showed that deamidation disrupts the interaction between UBC13 and TRAF6^RING^ by three mechanisms: i) A new intramolecular R14/E100 salt-bridge appears in dUBC13, which competes with a critical intermolecular salt-bridge in the native complex, ii) the salt-bridge competition also perturbs the UBC13/TRAF6^RING^ transient complexes to inhibit association, and iii) repulsion between the negatively charged E100 and the negatively charged interface of TRAF6^RING^ perturbs the transient complexes to reduce association between UBC13 and TRAF6^RING^. The effect of each mechanism on the binding was confirmed by binding studies using appropriate substitutions in either UBC13 or TRAF6^RING^. The impact of deamidation on transient interactions was also observed using another RING domain from RNF38, indicating that the mechanism could be ubiquitous for UBC13/RING complexes. Our study highlights that deamidation of residues that do not directly participate in the E2/E3 interaction but are close to the interface can effectively modulate the interaction by perturbing the native and transient intermolecular contacts. The mechanism of regulating protein-protein transient interactions by post-translational modifications could be at play in other quintessential signaling pathways.

## Results

### Deamidation abolishes the interaction between UBC13 and TRAF6^RING^

Two different constructs of TRAF6 was used in this study. The isolated RING domain TRAF6^RING^ (aa: 50-124) was the shorter construct, while the longer construct included the RING domain and three ZF domains (TRAF6^RZ3^, aa: 50-211). By the three dimensional structures, TRAF6^RING^ interacts with UBC13 and with the donor Ub in the UBC13~Ub conjugate, while the interaction of ZF domains with donor Ub further stabilizes the UBC13~Ub/TRAF6^RZ3^ complex (Middleton et al. 2017a). Both the E2/E3 complex of dUBC13/TRAF6^RZ3^ and dUBC13/TRAF6^RING^ had reduced polyubiquitination activity than the corresponding wild-type complexes (Figure 1A, 1B). Deamidation could have some allosteric effect at the active site or Ub-binding site, which may deactivate UBC13. However, a comparison of the extent of polyubiquitination by the UBC13/MMS2 heterodimer in the absence of E3 was similar between UBC13 and dUBC13, indicating that E2 activity is not altered upon deamidation (Figure 1C).

**Figure 1.**
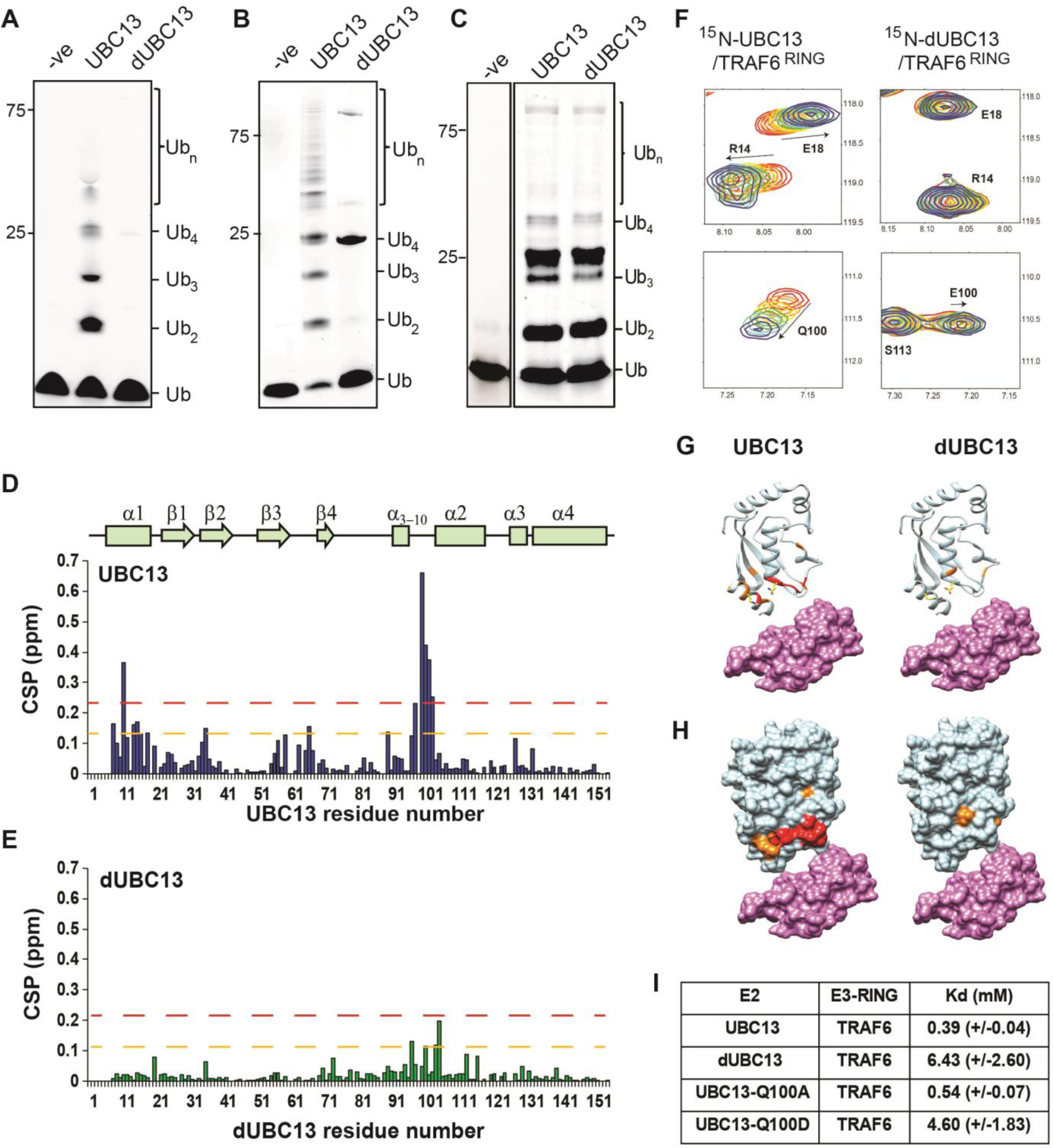
Interactions between UBC13 and TRAF6^RING^ studied by NMR. A) *In-vitro* ubiquitination reaction was carried out using TRAF6^RZ3^ as the E3 and UBC13 or dUBC13 as the E2 for 10 min. Mms2 was used as a co-factor in the reaction. The –ve lane is the same reaction without ATP. B) *In-vitro* ubiquitination reaction was carried out on GST beads for 10 min using GST-TRAF6^RING^ as the E3 and UBC13 or dUBC13 as the E2. Mms2 was used as a co-factor in the reaction. The –ve lane is the same reaction without ATP. B) *In-vitro* ubiquitination reaction carried out for 30 min using UBC13 or dUBC13 as the E2 and Mms2 as its co-factor. C)The CSPs for residues in UBC13 upon binding to TRAF6^RING^. The chemical shift perturbations (CSP) between the free and the bound form are calculated as CSP = [(δ^H^free – δ^H^bound)^2^+ ((δ^N^free – δ^N^bound))^2^]^1/2^, where δ^H^ and δ^N^ is the chemical shift of the amide hydrogen and nitrogen, respectively. The orange and red dashed lines correspond to Mean+SD and Mean+2*SD, respectively. The secondary structure alignment of UBC13 against its sequence is provided above the plot. D) The CSPs for residues in dUBC13 upon binding to TRAF6^RING^. The dashed lines are replicated from B). E) Two regions of the titration HSQC spectra are expanded to show UBC13, but not dUBC13 peaks shift upon titration with TRAF6^RING^. Significant CSPs were mapped on the UBC13 and dUBC13 structure both in the F) ribbon and G) surface representation. The UBC13 and dUBC13 are colored in light blue. The residues with CSPs above Mean+SD and Mean+2*SD are colored in orange and red, respectively. The surface of TRAF6^RING^ domain shown in magenta. The UBC13/TRAF6^RING^ complex is modeled from PDB 3HCU. H) The measured dissociation constants of UBC13 and its mutants with TRAF6^RING^ are provided as Mean+/− SD.

Deamidation could misfold UBC13 or inhibit the interaction between UBC13 and TRAF6^RING^. These possibilities were examined by studying the impact of deamidation on the structure of UBC13 and its interaction with TRAF6^RING^. ^15^N-labeled UBC13/dUBC13 and unlabelled TRAF6^RING^ were expressed in *E. Coli* and purified. Unlabelled TRAF6^RING^ was titrated into a sample of ^15^N-UBC13, and the binding was detected by ^15^N-edited Heteronuclear Single Quantum Coherence (HSQC) NMR experiments (Figure S1A). Perturbations due to the altered chemical environment upon ligand binding induce changes in the chemical shift of the backbone amide resonances. The chemical shift perturbations (CSP) plotted in Figure 1D shows that the significant perturbations in UBC13 occur in the α1-helix, and the loop between α_3-10_ and α2-helix, which is the canonical interface of UBC13/TRAF6^RING^ complex (Figure 1G, 1H). The resonance shifts can be plotted against the ligand: protein concentration, and fit to yield the K_d_ of interaction. The peak shifts in UBC13 titration spectra were fitted to yield a K_d_ of 0.39 (±0.04) mM (Figure 1I and S1B).

Backbone amide chemical shifts can report if the protein’s structure is affected by the substitution/modification of a residue. The backbone chemical shifts of UBC13 and dUBC13 resonances were assigned using ^13^C, ^15^N labeled samples, and standard triple resonance NMR experiments. The CSPs between UBC13 and dUBC13 indicated that apart from the residues immediately next to Q100 in sequence, only R14 and L15 of the N-terminal α1-helix are affected upon Q100E substitution (Figure S2A). The absence of major CSPs in the rest of dUBC13 indicated that deamidation did not change the fold of UBC13. Moreover, analysis of the backbone and C_β_ chemical shifts in dUBC13 confirmed that the secondary structure was unperturbed (Figure S2B). Unlabelled TRAF6^RING^ was then titrated into a sample of ^15^N-dUBC13, and the binding was detected by ^15^N-edited HSQC experiments (Figure S3). Negligible peak shifts were detected in dUBC13 even when the TRAF6^RING^ was titrated up to 6-fold higher than dUBC13, indicating that TRAF6^RING^ does not bind dUBC13 (Figure 1E-H, S3B). A rough estimate of the K_d_ was obtained by comparing the CSPs at the UBC13 interface at the same protein: ligand concentration between UBC13 and dUBC13, which suggested the K_d_ of 6.34 (±2.6) mM. If the reduced affinity is precisely due to the negative charge at residue 100, then substitution of Q100 with a neutral amino acid should not affect the affinity of the complex. TRAF6^RING^ was titrated to Q100A-UBC13, and the measured K_d_ was 0.54 (±0.07) mM (Figure 1I), which is similar to UBC13/TRAF6^RING^. However, when Q100 was substituted with another acidic residue aspartate (D), the affinity dropped significantly (Kd ~ 4.6 (±1.8)) mM, Figure 1I), confirming that substitution at Q100 with a negatively charged residue inhibited the interaction between UBC13 and TRAF6^RING^. Altogether, deamidation inhibited the binding of TRAF6^RING^ to UBC13, but neither changed the fold of UBC13 nor its intrinsic enzymatic activity.

### Deamidation triggers the formation of an intramolecular salt-bridge in UBC13

In the UBC13/TRAF6^RING^ native complex structure, Q100 does not form any contact with TRAF6^RING^ (Figure 2A). Hence, the mechanism underlying reduced binding due to deamidation of Q100 is nonintuitive. The surface electrostatic potentials calculated for UBC13 and TRAF6^RING^ indicated charge complementarity at the interface (Figure 3A). The UBC13 interface was positively charged, while the TRAF6^RING^ was negatively charged. Notably, Q100 lies in the vicinity of a network of electrostatic interactions involving helix-1 of UBC13 (R6 and R14) and the first zinc-coordination motif (D57 and E69) of the TRAF6^RING^ (Figure 2A). Possibly, the introduction of an additional negative charge upon deamidation could perturb the interfacial electrostatic contacts to disrupt the interaction.

**Figure 2:**
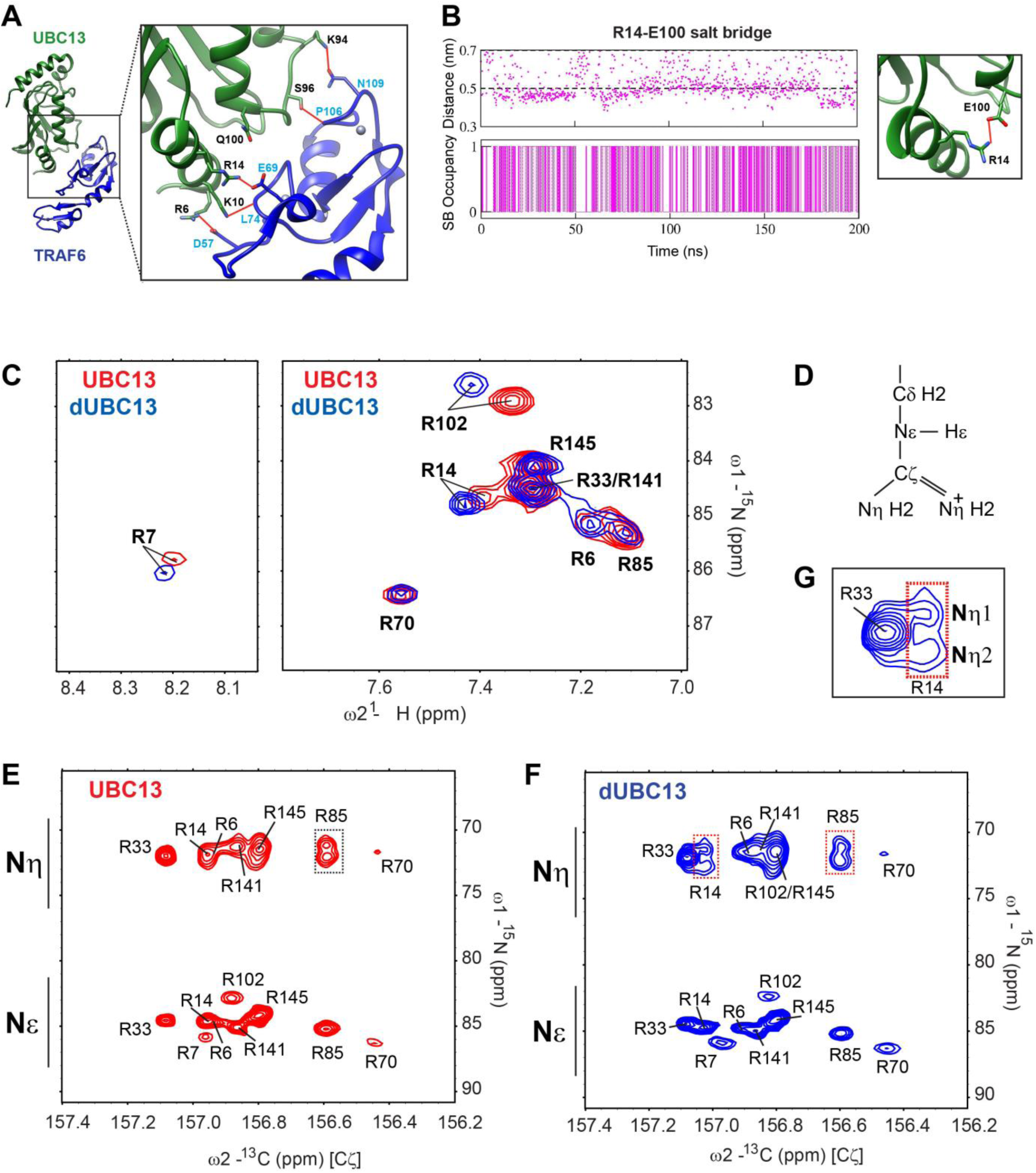
A new intramolecular salt-bridge forms upon deamidation of UBC13. A) Crystal structure of the UBC13/TRAF6^RING^ complex (PDB 3HCU, Left). Inset (right) indicates the position of Q100 and the network of salt-bridges/hydrogen bonds (red connecting lines) at the interface of UBC13/TRAF6^RING^ complex. B) The distance between R14-Cζ and E100-Cδ atoms against time in the conventional MD simulation of dUBC13 is given (top). The presence/absence of a salt-bridge based on a 0.5 nm cutoff value are digitized to a Markov chain, where the presence of a salt-bridge is 1 and absence is 0. C) Overlay of UBC13 and dUBC13 ^15^N-^1^H HSQC spectra zoomed around the Arginine Nε-Hε resonances, shows that R7, R14, and R102 sidechain resonances shift upon deamidation. D) Schematic of Arginine sidechain atoms. E) and F) are the ^15^Nε/η-^13^Cζ correlation spectra for UBC13 and dUBC13, respectively. The ^15^Nε/η and ^13^Cζ resonance shifts are in the x- and y-axis, respectively. The R33 and R14 ^15^Nη resonances of dUBC13 are expanded in G).

**Figure 3.**
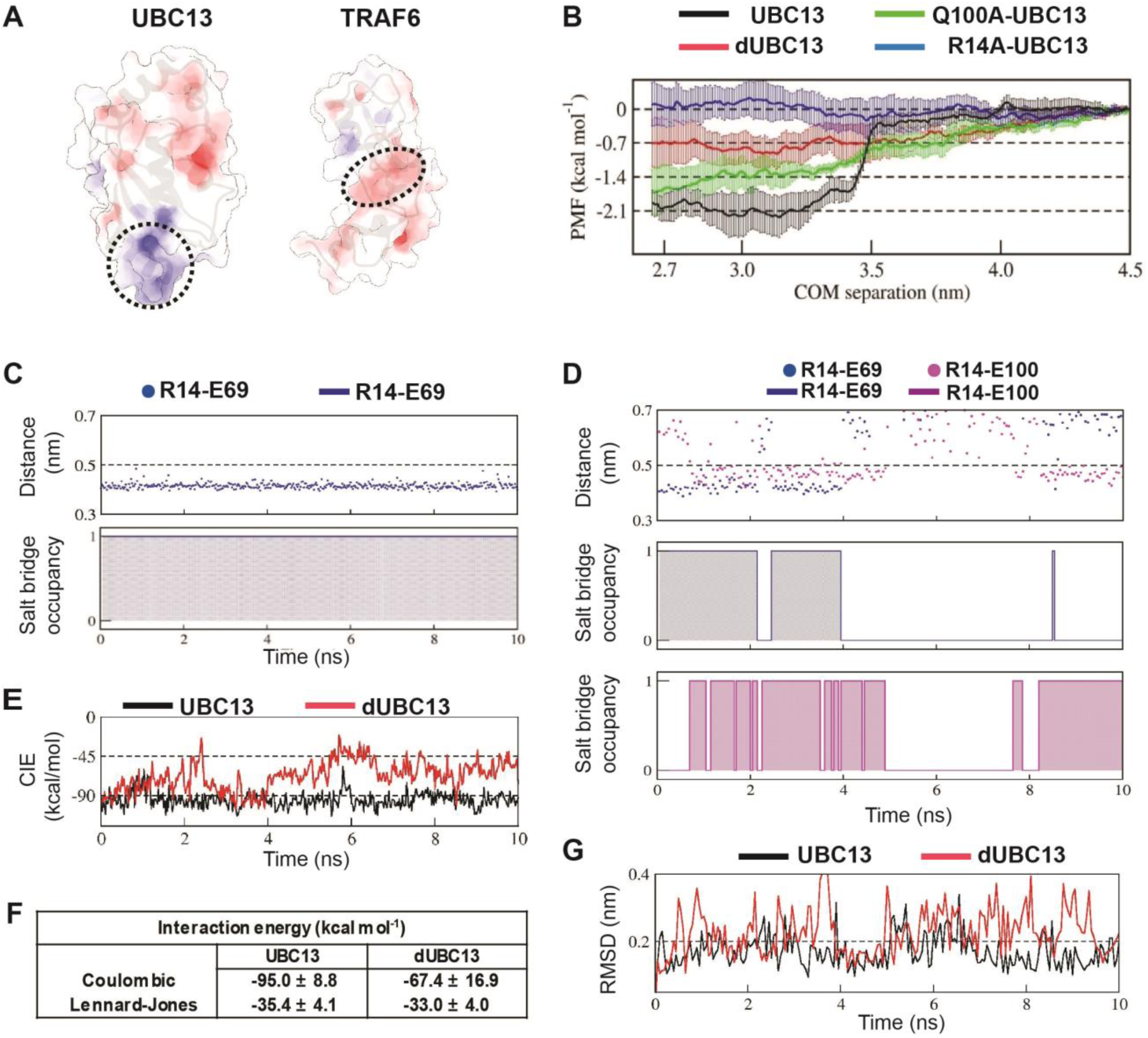
Deamidation induces salt-bridge competition to weaken UBC13/TRAF6^RING^ interaction. A) Surface electrostatic potentials of UBC13 and TRAF6^RING^. Circled regions indicate the complementary electrostatic surfaces on UBC13 and TRAF6^RING^ interfaces. The color scale ranges from −8 to +8 kT/e. B) Potential of mean force (PMF) profiles as a function of COM separation along the x-axis for the association between UBC13 variants and TRAF6^RING^. PMF profiles were calculated by averaging over five PMF profiles ranging from 2.5 ns to 10 ns. Error bars represent ± one standard error of the mean (SEM). C) Stability and occupancy of the R14-E69 salt-bridge in the native window for the wild-type complex. D) Stability and occupancy of the R14-E69 and R14-E100 salt-bridge in the native window for the dUBC13/ TRAF6^RING^ complex. In C) and D), distance plots indicate the distance between R14 Cζ and E69/E100 Cδ atoms. The salt-bridge occupancy plots were generated as in Figure 2B. E) Coulombic interaction energies (CIE) between UBC13 wild-type/Q100E and TRAF6^RING^ in native windows (mean COM separation = 2.7 nm) from the US simulations. The table in (F) reports the Mean ± SD of the interaction energies over 10 ns. G) RMSD of TRAF6^RING^ (aa:70-109) against time in the UBC13 and dUBC13 complexes.

A conventional MD simulation (200 ns) of free dUBC13 was performed to investigate if deamidation alters any electrostatic interactions within UBC13. Interestingly, an intramolecular salt-bridge was observed between R14 and E100 in dUBC13 (Figure 2B), which correlates well with the backbone chemical shifts observed at R14 upon deamidation (Figure S1B). The formation of the new salt-bridge involving R14 and E100 in dUBC13 was further investigated by NMR spectroscopy. ^15^N-edited HSQCs of the Arginine Nε-Hε groups were collected for both UBC13 and dUBC13, which showed shifted resonances for the arginine sidechains (R7, R14, and R102) around residue E100, indicating that these sidechain conformations are perturbed (Figure 2C). Unfortunately, the ^1^Hη atoms are invisible in the NMR spectra due to chemical exchange with the solvent at physiological pH, and millisecond timescale rotations around the Cζ-Nη/ Cζ-Nε bonds (Figure 2D). Hence, detection of arginine sidechain-mediated interactions (hydrogen bonds and salt-bridges) through ^1^Hη atoms is difficult. However, such interactions can be inferred from the reduced mobility of their sidechains observed in ^15^Nη/ε(F1)-^13^Cζ(F2) correlation spectra (Yoshimura et al. 2017). For example, R85 forms salt-bridge with D81 in UBC13, which reduces the sidechain mobility of R85 sidechain and gives rise to two separate resonances for R85 Nη atoms in the ^15^Nε/η-^13^Cζ correlation spectra (Figure 2E). The side-chains of free arginines rotate faster and their corresponding Nη atoms have a single averaged resonance in the spectra (Figure 2E). The R14 Nη atoms had two separate resonances in dUBC13 but not UBC13 (Figure 2F, expanded in 2G), implying that R14 forms a salt-bridge in dUBC13, consistent with the MD simulations. The salt-bridge persisted in the Q100D-UBC13 (Figure S4), correlating well with its reduced affinity for TRAF6^RING^.

### Deamidation triggers salt-bridge competition at the UBC13-TRAF6^RING^ interface

At the interface of UBC13/TRAF6^RING^, R14 of UBC13 forms an intermolecular salt-bridge with E69 of TRAF6^RING^ (Figure 2A). MD simulations of dUBC13 suggested that the deamidation-induced intramolecular R14/E100 salt-bridge may interfere with the intermolecular R14/E69 salt-bridge to destabilize the complex. Conventional MD simulations were performed with both UBC13/TRAF6^RING^ and dUBC13/TRAF6^RING^ to test this hypothesis. The stability of the complex during the simulation was determined by the rmsd of TRAF6^RING^ with respect to the crystal structure. Higher rmsd implied lower stability of the complex. The UBC13/TRAF6^RING^ complex was stable throughout (Figure S5A). The dUBC13/TRAF6^RING^ complex was unstable for a significant duration in a trajectory, wherein the rmsd of TRAF6^RING^ increased substantially (Figure S5B).

Five interfacial contacts were chosen as the reporter of the interaction between UBC13 and TRAF6^RING^ (Figures S5). The three selected electrostatic contacts were the R14/E69 salt-bridge, R6/D57 salt-bridge, and S96/P106 hydrogen bond (Figure 2A and S5A). The two selected hydrophobic interactions were R7/I72 and M64/L74 (Figure S5D). The other interfacial contacts were mostly unstable in simulations and were omitted from our analysis (Figure S5G and S5H). No major instability was observed in the hydrophobic contacts between UBC13 and dUBC13 (Figure S5E and S5F). However, concurrent to the high rmsd in dUBC13/TRAF6^RING^ complex, the R14/E69 intermolecular salt-bridge disrupted and the R14/E100 intramolecular salt-bridge formed, suggesting a role of the salt-bridge competition in destabilizing the native complex (Figure S5B).

The Umbrella Sampling (US) method was used to determine the change in binding free energy of the UBC13/TRAF6^RING^ complex upon deamidation. The US method quantifies the amount of work done to form the native complex from a distant separation, where the two proteins are non-interacting. The potential of mean force (PMF) for the association between the proteins is computed as a function of their center of mass (COM) separation. As a control of the protocol, US simulations were performed to study the association between Barnase and Barstar (Figure S6, Table 1 and Table S1). The observed PMF profile and ΔG_PMF_ (−12.6 kcal mol^−1^) of the Barnase/Barstar complex are consistent with the high-affinity of the complex (K_d_~136 fM) and previous MD studies (Hoefling & Gottschalk 2010; Wang et al. 2010).

**Table 1.** Association PMF determined by Umbrella sampling MD.

| Complex | $\langle \Delta G_{\text{PMF}} \rangle \text{ (kcal mol}^{-1}\text{)}$ |
| --- | --- |
| Barstar/Barnase | $-12.60 (\pm 0.76)$ |
| UBC13/TRAF6 | $-2.03 (\pm 0.27)$ |
| dUBC13/TRAF6 | $-0.67 (\pm 0.50)$ |
| Q100A-UBC13/TRAF6 | $-1.64 (\pm 0.49)$ |
| R14A-UBC13/TRAF6 | $+0.04 (\pm 0.38)$ |

\*Standard error of mean is indicated in brackets

Destabilizing modifications/substitutions in UBC13 will increase the overall work done (ΔG_PMF_) to form the native complex. The US simulations were carried out for both the UBC13/TRAF6^RING^ and dUBC13/TRAF6^RING^ complexes (Figure 3B, Table 1, and Table S1). At the 3.2 nm COM separation, the PMF profile of UBC13/TRAF6^RING^ decreased sharply by 2.0 kcal mol^−1^ and subsequently plateaued (Figure 3B). In contrast, ΔG_PMF_ of dUBC13/TRAF6^RING^ gradually reduced by only 0.7 kcal mol^−1^, suggesting that deamidation reduces the binding energy between UBC13 and TRAF6^RING^. The difference in ΔG_PMF_ between the deamidated and wild-type (wt) complex was in good agreement with the decrease in binding energy observed in the NMR titrations (ΔΔG_PMF_ =1.4 kcal mol^−1^, ΔΔG_binding_=1.6 kcal mol^−1^). The ΔG_PMF_ of Q100A-UBC13/TRAF6^RING^ is, however, similar to the wt complex (Table 1).

The stability of the intermolecular contacts could be compared between the two complexes within the native window in the US simulations. While the R14/E69 salt-bridge was stable throughout in UBC13/TRAF6^RING^, it disrupted within 4ns due to salt-bridge competition in dUBC13/TRAF6^RING^ (Figure 3C, 3D). The instability of R14/E69 salt-bridge correlates well with lower Coulombic Interaction Energy (CIE) and higher rmsd of TRAF6^RING^ (Figure 3E and 3G). A significant difference of ~30 kcal mol^−1^ was observed in the mean CIE between the two complexes (Figure 3F). In contrast, the difference in Lennard-Jones interaction energy was mere ~2.4 kcal mol^−1^, suggesting that deamidation primarily inhibits the electrostatic interactions. Moreover, the PMF profile of R14A-UBC13/TRAF6^RING^ was flat with ΔG_PMF_~0 kcal mol^−1^, suggesting that the interactions mediated by R14 are essential for the complex (Figure 3B). Overall, the native complex simulations and US simulations indicate that salt-bridge competition reduces the binding energy of the dUBC13/TRAF6^RING^ complex.

### Salt-bridge competition enhances the dissociation of dUBC13/TRAF6^RING^

To capture the effects of deamidation on the dissociation of the native complex, steered MD (SMD) simulations were performed for both the wt and dUBC13 complex. Ten independent, 30 ns long SMD simulations were performed, wherein the COM of the TRAF6^RING^ was separated from UBC13 by ~6 nm at a slow rate of 0.125 nm ns^−1^. The COM position of UBC13 was kept fixed. The average maximum unbinding force (F_max_) and work (W) profiles were determined by combining all SMD simulations (Figure 4A, B and Table 2). The distribution of F_max_ and W for individual SMD trajectories are given in Figure 4C and Table S2. Compared to the UBC13/TRAF6^RING^ complex, lesser F_max_ and W were required to dissociate the dUBC13/TRAF6^RING^ complex, indicating reduced stability due to deamidation of UBC13. The mean occupancy of the R14/E69 salt-bridge was lower for dUBC13/TRAF6^RING^ compared to UBC13/TRAF6^RING^ (Figure 4D, S7, S8), because the salt-bridge competition disrupted the R14/E69 salt-bridge for prolonged periods in several trajectories (Figure S8 - trajectories 3,4,5 & 9). As a result, the W of these trajectories were lower than the minimum value of W (28 kcal.mol^−1^) observed in any wt trajectory (Table S2). A similar destabilization of the complex was observed when the SMD was repeated with R14A-UBC13, where the R14/E69 salt-bridge is absent. Overall, R14/E69 salt-bridge was unstable in the dUBC13/TRAF6^RING^ complex, which reduced the unbinding work required to dissociate the complex.

**Figure 4.**
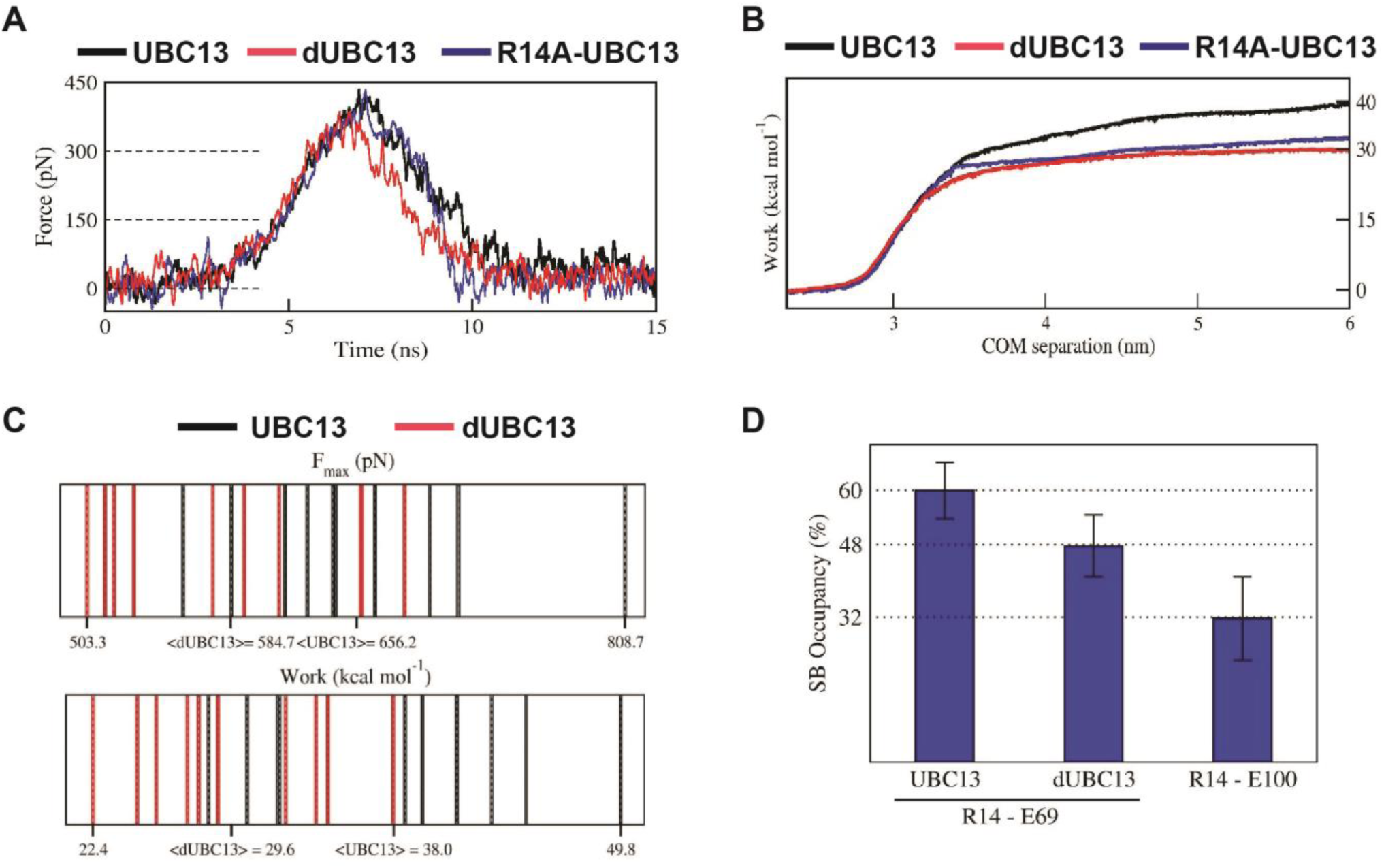
Steered MD of the UBC13/TRAF6^RING^ and dUBC13/TRAF6^RING^ complex. A) Average force-extension profiles for wild-type and mutant complexes are plotted. The profiles were smoothened over 250 ps time intervals. B) The plot of average work against COM separation indicated the cumulative work done to separate wild-type and mutant complexes. C) Distribution of F_max_ (top) and unbinding work (bottom) values obtained from ten individual SMD trajectories for wild-type (black) and mutant (red) complexes are shown. In each plot, the minimum, maximum, and average (< >) values are indicated on the x-axis. Both plots reveal a shift in the range of F_max_ and unbinding work towards lower values for the dUBC13/TRAF6^RING^ complex, which correlate with the reduced binding. D) Mean ± one standard error (SEM) of R14-E69/E100 salt-bridge (SB) occupancies averaged from the first 15 ns of ten individual SMD trajectories are shown. SEM for all three occupancies is within 10%.

**Table 2.** F_max_ and unbinding work determined by Steered MD.

| Complex | $\langle F_{\text{Max}} \rangle$ (pN) | $\langle \text{Work} \rangle$ (kcal mol<sup>-1</sup>) |
| --- | --- | --- |
| UBC13/TRAF6 | 466.7 ($\pm 31.5$) | 38.1 ($\pm 2.2$) |
| dUBC13/TRAF6 | 425.9 ($\pm 42.2$) | 29.6 ($\pm 1.6$) |
| R14A-UBC13/TRAF6 | 450.4 ($\pm 45.9$) | 31.9 ($\pm 2.1$) |

\*Standard error of mean is indicated in brackets

The order in which the contacts disrupted during dissociation was compared in a typical SMD trajectory for both complexes (Figure 5A and Movies S2, S3). Whereas the other significant contacts disrupted within 8 ns, the R14/E69 salt-bridge persisted till the complex dissociated completely at 15 ns (Figures 5A-5C and S9). Given the R14/E69 salt-bridge was stable in both conventional and SMD simulations, it could be critical for the stability of the wt complex. On the contrary, R14/E69 salt-bridge disrupted within 3 ns in the dUBC13/TRAF6^RING^ complex, due to competition from the R14/E100 salt-bridge (Figures 5D-F). The R14/E100 salt-bridge in dUBC13 remained stable after dissociation of TRAF6^RING^ (Figure 5E). Collectively, the SMD suggested that R14/E69 formed a critical interfacial salt-bridge, and competition from E100 disrupted the salt-bridge to enhance dissociation of the complex.

**Figure 5.**
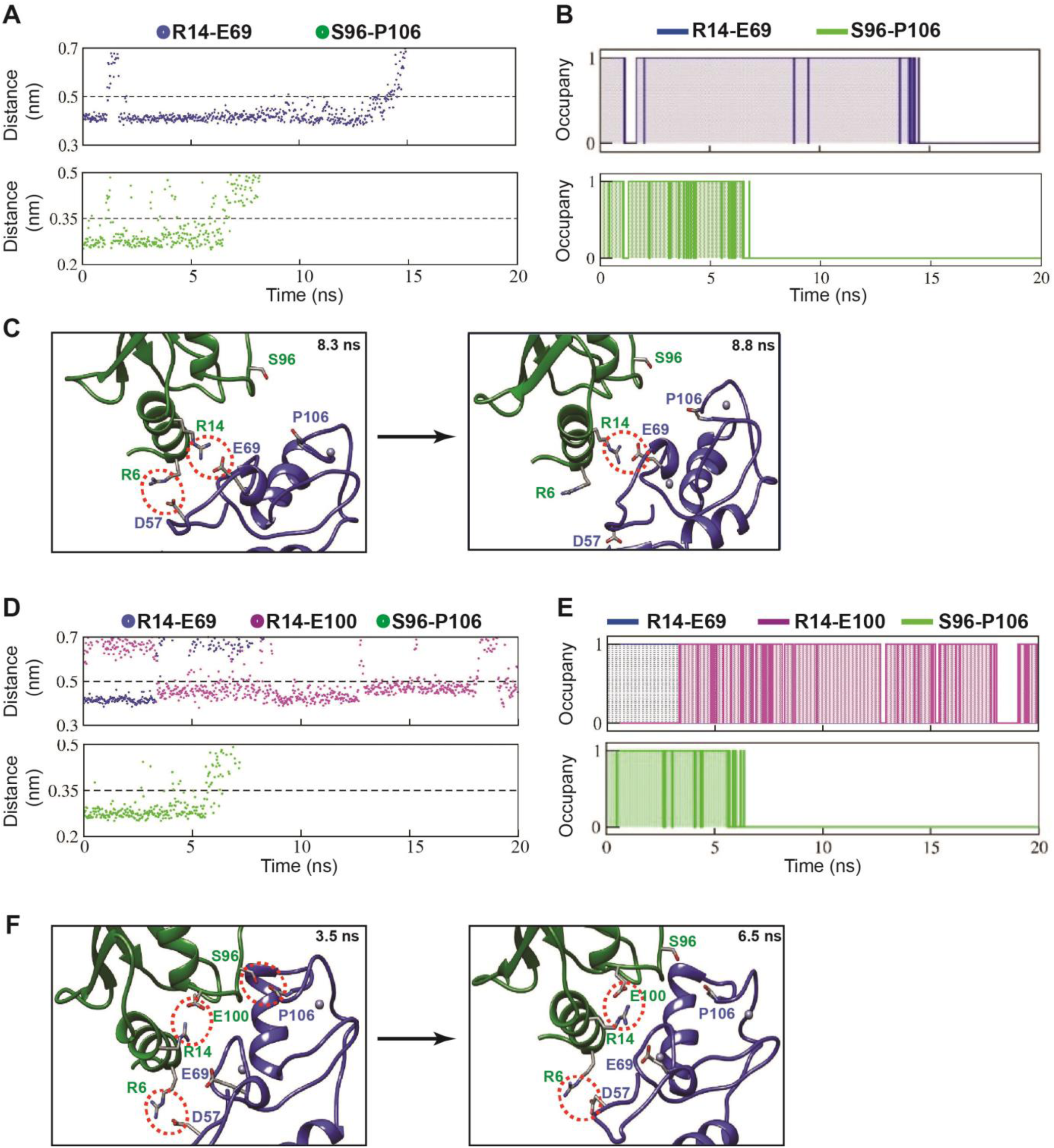
Pathway of UBC13/TRAF6^RING^ complex dissociation from steered MD. A) Top: The distance between R14-Cζ and E69-Cδ atoms against time during SMD. Bottom: The distance between S96-Oγ and P106-O atoms against time during SMD. B) Occupancy plots of A) calculated as in Figure 2B. C) Two snapshots from the trajectory in A) are shown. S96-P106 contact is disrupted at 8.3 ns followed by the R6-D57 salt-bridge break at 8.8 ns. Red dotted circles indicate the polar contacts. D) Same as A) for the dUBC13/TRAF6^RING^ complex. The distance between R14-Cζ and E100-Cδ atoms against time is added here. E) Occupancy plots of R14-E69 (blue), R14-E100 (magenta) and S96-P106 (green) contacts for the dUBC13/TRAF6^RING^ complex. F) Two snapshots from trajectory analyzed in D) showed that competition between R14-E100/E69 salt-bridges as the complex starts to dissociate at 6.5 ns.

### Repulsive interactions destabilize the dUBC13/TRAF6^RING^ transient complex ensemble

A native protein complex forms through an intermediate species referred to as the transient/encounter complex ensemble (Schreiber, Haran & Zhou 2009). These complexes are formed by native/non-native long-range electrostatic interactions and have greater oriental freedom compared to the native state (Tang, Iwahara & Clore 2006). Since charge distribution is altered upon deamidation, its effect on the complex was estimated based on the transient complex theory (Alsallaq & Zhou 2008). The method uses the native complex structure and computes association rates (k_on_) from a basal association rate constant (k_a0_) due to random diffusion and the electrostatic interaction free energy of the transient complex (ΔG_el_). The k_a0_ values, ΔG_el_ values, and k_on_ rates computed for two complexes are summarised in Table 3. The k_on_ of UBC13/TRAF6^RING^ decreased by five-fold at high salt compared to low salt, confirming that electrostatic interactions are critical for the transient complex. The k_on_ of dUBC13/TRAF6^RING^ reduced by three-fold compared to the wt complex at low salt. However, at high salt, the drop in k_on_ was only 1.6-fold, indicating that deamidation modulates the electrostatic interactions. Since the electrostatic interactions were partially screened at high salt, the effect of deamidation attenuated. The position of individual atoms within each input structure remained fixed in transient-complex theory calculations, and hence, the competition between salt-bridges was absent during these calculations. Therefore, the changes in k_on_ upon deamidation were solely due to electrostatic repulsion between E100 of UBC13 and the acidic residues on TRAF6^RING^.

**Table 3.** Electrostatic free energies and association rate constants (k_a0_/k_on_) of UBC13/TRAF6^RING^ calculated using TransComp web server.

| Transient complex Ensemble | $k_{a0}$<br>($10^6 \cdot M^{-1} s^{-1}$) | 10 mM NaCl | | 100 mM NaCl | |
| --- | --- | --- | --- | --- | --- |
| | | $\Delta G_{el}$<br>(kcal mol<sup>-1</sup>) | $k_{on}$<br>($10^6 \cdot M^{-1} s^{-1}$) | $\Delta G_{el}$<br>(kcal mol<sup>-1</sup>) | $k_{on}$<br>($10^6 \cdot M^{-1} s^{-1}$) |
| UBC13/TRAFF6 | 0.57 | -2.32 | 28.7 | -1.33 | 5.43 |
| dUBC13/TRAFF6 | 0.63 | -1.59 | 9.36 | -0.99 | 3.43 |
| R14A-UBC13/TRAFF6 | 0.62 | -1.16 | 4.42 | -0.62 | 1.78 |

The US simulations were analyzed to obtain molecular details of the effect of deamidation on transient complexes (Figure 6). A suitable point to observe the transient complexes is the US window corresponding to 3.0 nm COM separation (Figure 3B). A prominent transient interaction observed in the UBC13/TRAF6^RING^ complex was between R14 in UBC13 and D57 in TRAF6, which prompted the association between UBC13 and TRAF6^RING^. Subsequently, other contacts formed to stabilize the complex such that the distance between UBC13 and TRAF6^RING^ was invariant (Figure 6A-C). The CIE was favorable throughout the trajectory (Figure 6D). The R14/D57 interaction was, however, unstable in the case of dUBC13/TRAF6^RING^ (Figure 6C). Consequently, dUBC13 separated completely from TRAF6^RING^ within 2 ns, and the CIE dropped to zero (Figure 6B-D). Interestingly, in the first 2 ns, R14 and E100 simultaneously interacted with D57 (Figure 6E, 6F). While the R14/D57 interaction was attractive, the E100/D57 interaction was repulsive. The repulsive interaction destabilized the transient complex (higher CIE in Figure 6D) and triggered its dissociation. The R14/E100 intramolecular salt-bridge formed soon after dissociation, which would prevent further transient association between R14 and TRAF6^RING^ residues. Overall, the electrostatic repulsion between E100 and negatively charged interface of TRAF6^RING^ destabilizes the dUBC13/TRAF6^RING^ transient complexes.

**Figure 6.**
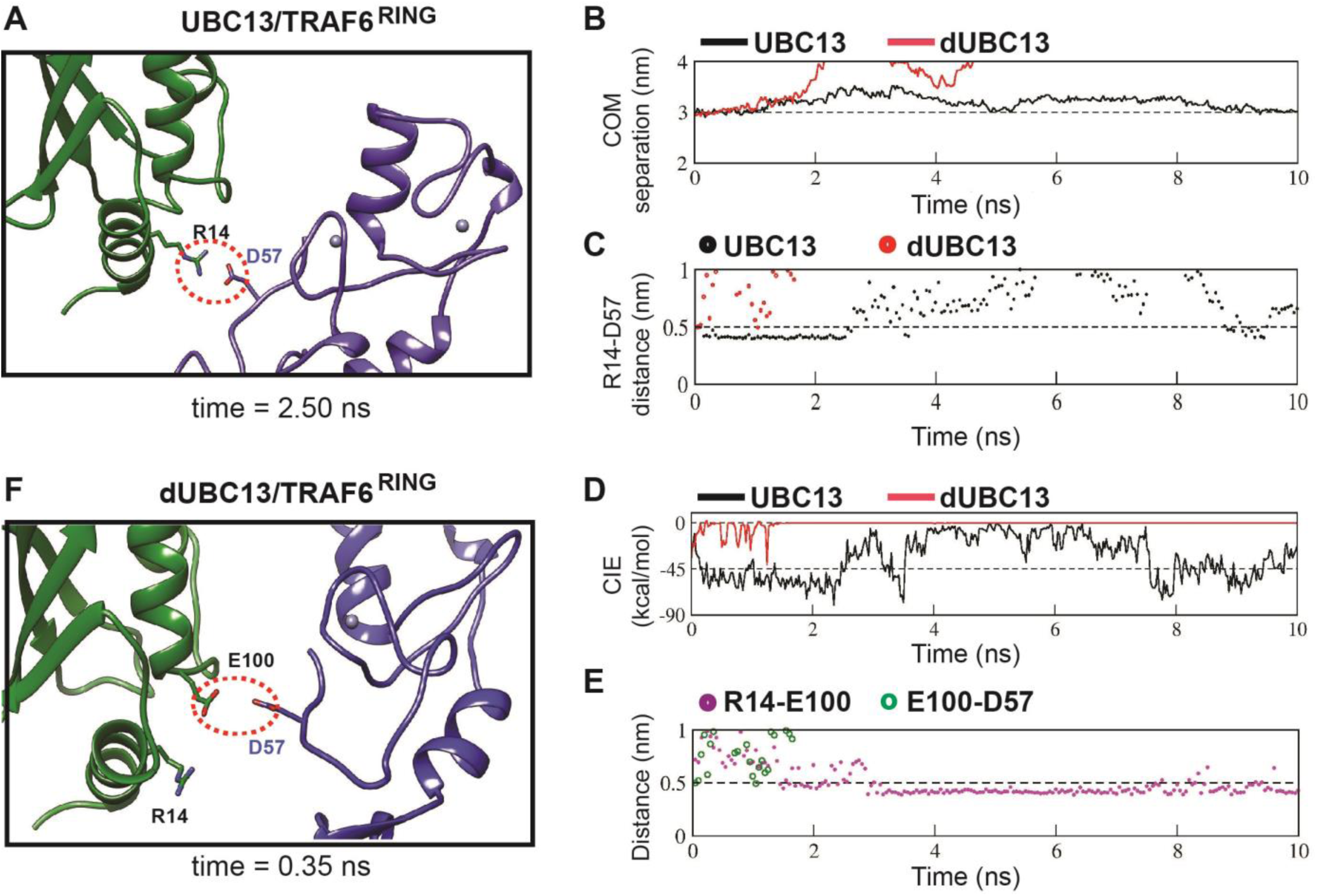
Analysis of transient complexes formed in the US simulations. The US window corresponding to 3 nm COM distance of separation was analyzed. (A) The transient complex of UBC13 and TRAF6^RING^ at t=2.50 ns is shown. A transient intermolecular salt-bridge between R14 and D57 is indicated by a red dotted circle. (B) The plot of COM separation against time for the UBC13/TRAF6^RING^ and dUBC13/TRAF6^RING^ complexes. (C) The distance between the R14-D57 transient contact is compared between the two complexes. The distances were measured between R14-Cζ and D57-Cγ atoms. (D) Coulombic interaction energy (CIE) plotted against time. (E) The contact distance of E100-D57 and R14-E100 contacts are shown against time for the dUBC13/TRAF6^RING^ complex. The distances were measured between R14-Cζ, E100-Cδ, and D57-Cγ atoms. (F) The transient complex of dUBC13 and TRAF6 at t=0.35 ns, where E100 contacted D57.

### Salt-bridge competition destabilizes the UBC13/TRAF6^RING^ transient complex ensemble

When the transient complex theory calculations were repeated for the R14A-UBC13/TRAF6^RING^ complex, the predicted k_on_ dropped significantly by six-fold at low salt (Table 3). Since repulsion due to E100 is absent in R14A-UBC13, the drop in k_on_ suggests that R14-mediated interactions are crucial for the transient complex. Molecular details of transient complexes observed in US simulations also suggested that R14 mediated interactions favor the association of UBC13/TRAF6^RING^ complex. Unbiased-association simulations were performed on the two complexes to confirm this observation. From these simulations, the association probability and stability of native-like transient complexes were assessed (Table 4). The mean time of association of native-like transient complexes for UBC13/TRAF6^RING^ was 3-fold higher at low salt than high salt, confirming that indeed the transient complex association was dependent on the long-range electrostatic interactions. However, the mean-time of association in dUBC13/TRAF6^RING^ was indifferent to differences in salt-concentrations, indicating that the critical transient electrostatic interactions are disrupted upon deamidation. The mean time of association dropped by 2.3 fold between UBC13/TRAF6^RING^ and dUBC13/TRAF6^RING^ at low salt, suggesting that deamidation affects the association of the native-like transient complex.

**Table 4.** Summary of unbiased-association MD simulations.

|  | Association Trajectories* (%) |  |  |  |
| --- | --- | --- | --- | --- |
|  | 10 mM NaCl<br>(low salt) |  | 100 mM NaCl<br>(high salt) |  |
|  | UBC13 | dUBC13 | UBC13 | dUBC13 |
| Native-like association | 53 (8) | 20 (3) | 27 (4) | 33 (5) |
| Non-native association | 47 (7) | 60 (9) | 40 (6) | 47 (7) |
| No association | 0 (0) | 20 (3) | 33 (5) | 20 (3) |
|  | Mean percentage time of native-like association (%)** |  |  |  |
|  | 32 (9) | 14 (9) | 11 (7) | 11 (5) |

\*The number of trajectories used is indicated in brackets.
\*\*Standard error of mean is shown in brackets.

The ensemble of transient complexes observed in all the unbiased association trajectories at low salt was compared between UBC13 and dUBC13 (Figure 7). The distribution of TRAF6^RING^ COMs around the helix 1 of UBC13 indicates that the TRAF6^RING^ positions are tightly clustered around the UBC13 interface (Figure 7A). The projections of the transient complexes on the coordinate planes showed that a cluster of the transient complexes coincided with the native complex. The position of TRAF6^RING^ is more dispersed when it interacts with dUBC13, and none of the clusters coincide with the native complex (Figure 7B). The transient complexes were clustered by pairwise rmsd, and the clusters with more than ten structures were plotted for UBC13/TRAF6^RING^ and dUBC13/TRAF6^RING^, in Figure 7C and 7D, respectively. Several clusters around R14 were absent in dUBC13/TRAF6^RING^ transient complexes when compared to the wt transient complexes, indicating that the R14-mediated long-range transient interactions are inhibited in dUBC13.

**Figure 7.**
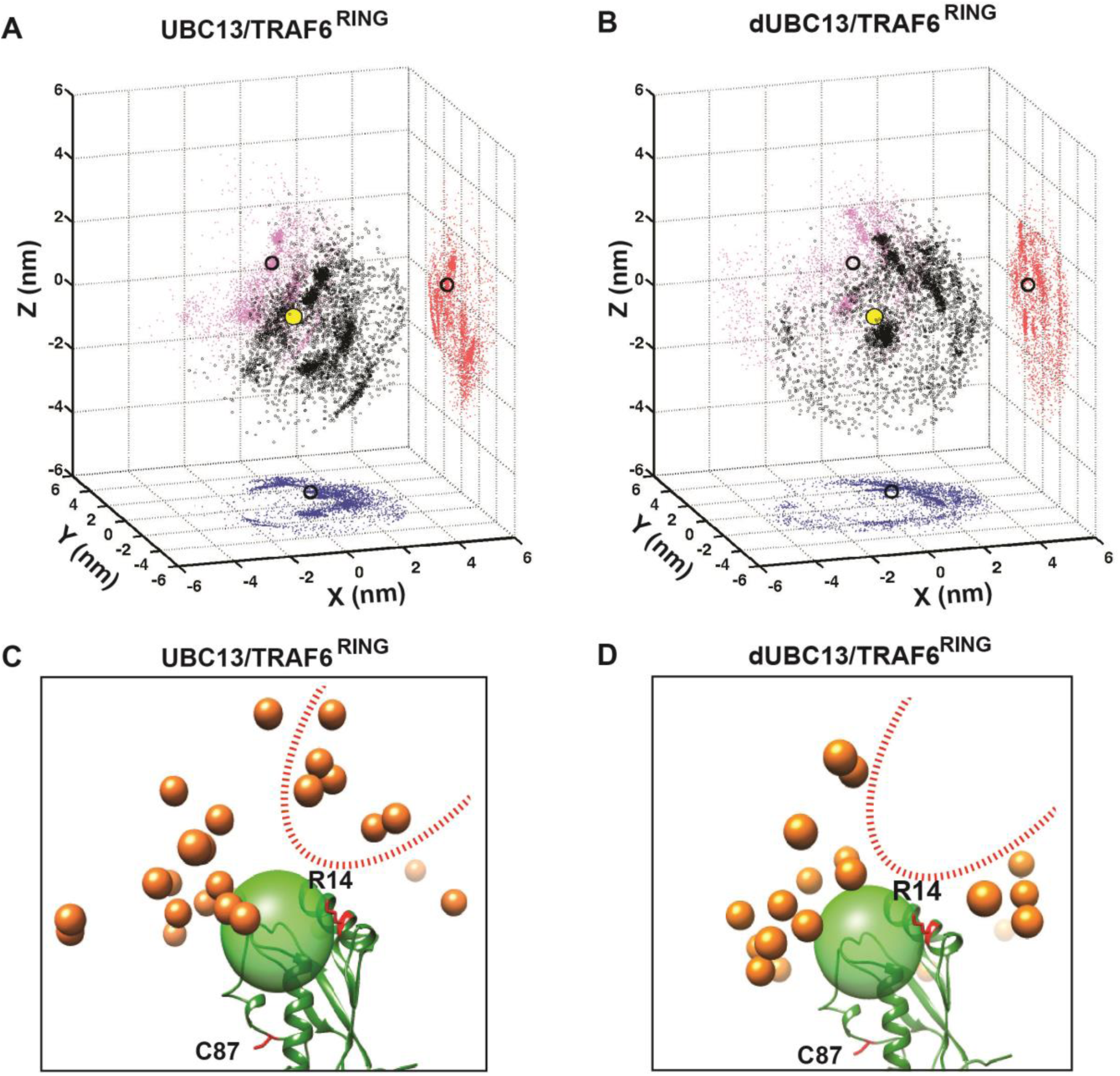
Transient complexes observed during unbiased association simulations. (A) Black open circles denote TRAF6^RING^ (aa:70-109) centers of mass around UBC13 in the Cartesian coordinate space. Red, pink and blue colored symbols denote projections of the centers of masses on the coordinate planes. Projections of the centers of mass of TRAF6^RING^ in the native complex are shown as black open circles on the coordinate planes. The center of mass of the UBC13 helix α1, which is at the center of UBC13 interface, is shown as a yellow sphere at the origin. (A) Same as in (B) for dUBC13. (C) TRAF6^RING^ clusters calculated based on pairwise RMSD between UBC13/TRAF6^RING^ complexes (cutoff = 0.45 nm) are shown as orange spheres. Only representatives for clusters with greater than ten structures are shown. The UBC13 is represented in ribbon and colored green. The center of mass of UBC13 interface is shown as a green transparent sphere with 1 nm radius. The sidechain of R14 is shown and colored red. (D) Same as (C) for dUBC13. The region near R14, where TRAF6^RING^ clusters were absent in dUBC13 is shown as a red dotted curve in (C) and (D).

The energy landscape correlating the free-energy with rmsd of TRAF6^RING^ and the minimum distance between interfaces shows that multiple low-energy transient intermediates are present in the UBC13/TRAF6^RING^ association pathway, which converged to the native complex (Figure S10A). In contrast, a few transient intermediates are present in dUBC13/TRAF6^RING^, and none exist in the native-like conformation (Figure S10B). The association trajectories were further analyzed to detect native-like complexes, whose rmsd to the native UBC13/TRAF6^RING^ structure is below 0.5 nm (Figure S11A). The analysis identified one trajectory at 10 mM NaCl, where the native complex formed for 50 ns during the simulations (Figure 8A, S11A). In this trajectory, TRAF6^RING^ associated with UBC13 within the first 10 ns via the R6/D57 and R14/E69 salt-bridges (Figures 8B, Movie S4). Consequently, the other relevant contacts formed, leading to the native complex at 25 ns, which persisted for the next 50 ns (Figure 8A, S12A). No native complex was detected in the dUBC13/TRAF6^RING^ trajectories at low salt conditions (Figures S11B). At higher salt, where electrostatic interactions are partially screened, a native-like complex was momentarily observed (Figures S11B, 8C, Movie S5). Herein, the initial association occurred through a formation of a non-native R6/E69 salt-bridge and the S96/P106 hydrogen bond at ~14 ns. Soon the R14/E100 salt-bridge competed to prevent the formation of R14/E69 salt-bridge, and the complex dissociated thereafter (Figures 8D and S12). In summation, unbiased association simulations indicate that the competition between salt-bridges and repulsive interactions perturb the ensemble of transient complexes to inhibit the association between dUBC13 and TRAF6^RING^.

**Figure 8.**
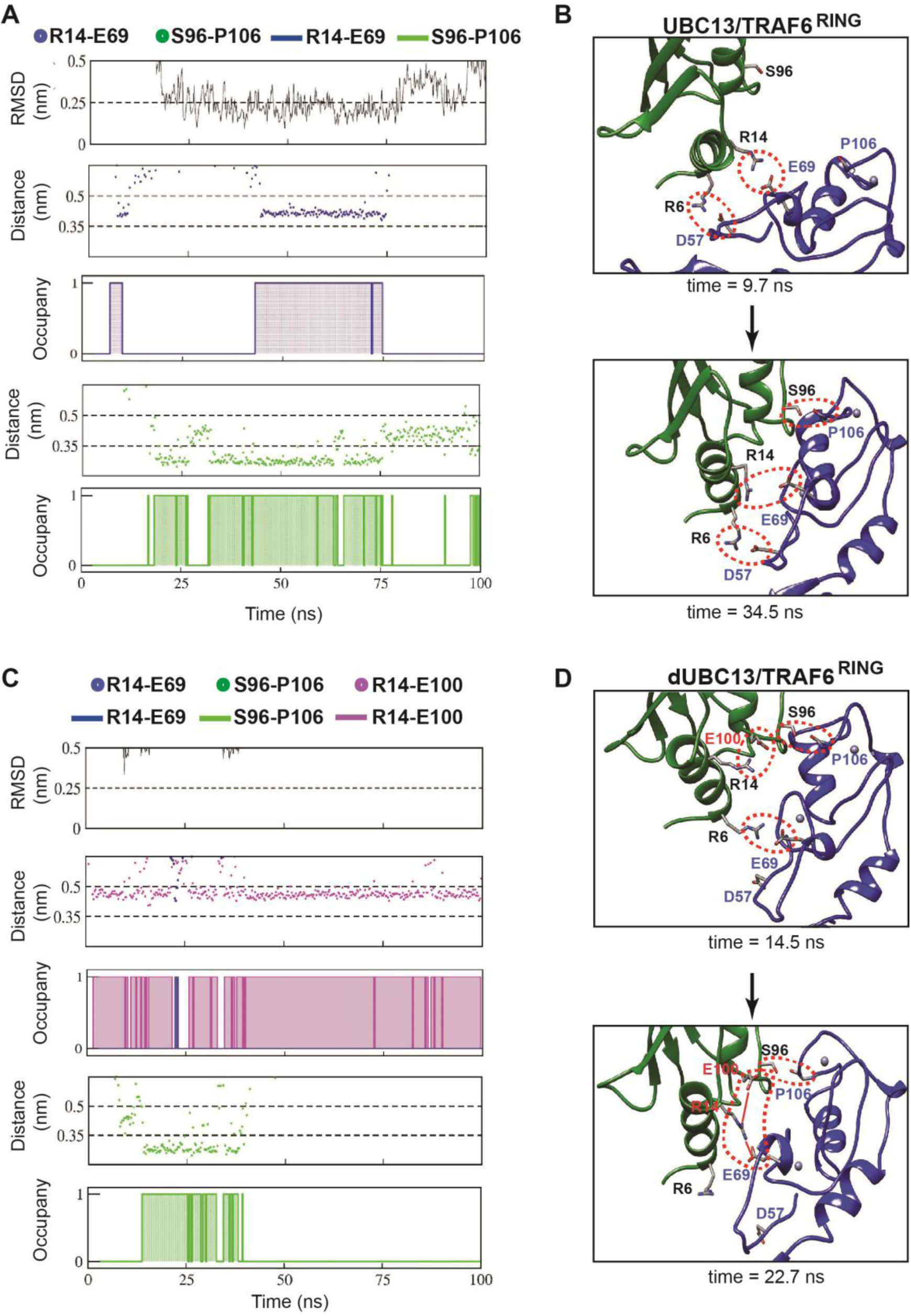
Association pathway of the UBC13/TRAF6 complex from unbiased MD trajectory. A) and C) shows the RMSD of TRAF6^RING^, the distance between polar contacts and occupancies of polar contacts in UBC13/TRAF6^RING^ and dUBC13/TRAF6^RING^ trajectory, respectively. The occupancies are calculated as in 2B. B) Two snapshots from a trajectory in A) is shown, where the initial contacts formed at 9.7 ns led to the native complex at 34.5 ns. The polar contacts at the interface are shown by red dotted circles. D) Two snapshots from a trajectory in B) is shown, where the transient contacts formed at 14.5 ns, but due to competition between the salt-bridges at 22.7 ns (shown by red lines), the complex dissociated subsequently.

### Binding studies delineate the effects of deamidation on the UBC13/TRAF6^RING^ complex

The MD simulations identified that deamidation destabilized the native complex by salt-bridge competition. In addition, the transient complexes were also destabilized by salt-bridge competition and repulsive interactions. NMR titrations were carried out using variants of UBC13 and TRAF6^RING^ to validate the effect of each mechanism on the affinity of the complex. E69 in TRAF6^RING^ was substituted to alanine to remove the acceptor of the R14/E69 intermolecular salt-bridge. The substitution mimics the effect of salt-bridge competition exclusively in the native complex. When ^15^N-UBC13 was titrated with E69A-TRAF6^RING^, the affinity dropped by 4-fold compared to the wt-complex (Figure 9A). Next, R14 in UBC13 was substituted with alanine. Such a substitution mimics the effect of salt-bridge competition in both the native and the transient complexes. When ^15^N-R14A-UBC13 was titrated with TRAF6^RING^, the affinity decreased by 11-fold (Figure 9A). Assuming the effect of salt-bridge competition on the transient and native complexes are mutually exclusive, the decrease in the affinity of the R14A-UBC13/TRAF6^RING^ complex would be a product of these two effects. Given the effect of salt-bridge competition in the native complex was 4-fold, the effect of salt-bridge competition on the transient complex would be 2.8-fold (=11/4). Similarly, the repulsive effect was assumed to be mutually exclusive to the salt-bridge competition. The combined effect of salt-bridge competition (both on the native and transient complex) and repulsion on the binding affinity is 17-fold, whereas the effect of the salt-bridge competition is 11-fold. Hence, the repulsive effect on the transient complex reduces the affinity by 1.5 fold (=17/11). The reduction in binding free energy due to repulsion is 0.24 kcal mol^−1^, which compares well with the ΔΔG_el_ = ΔG_el_^UBC13^ - ΔG_el_^dUBC13^ = 0.34 kcal mol^−1^ calculated by transient-complex theory at 100 mM NaCl (Table 3). Overall, titrations studies indicated that all three events have distinct effects on the binding between dUBC13 and TRAF6^RING^.

**Figure 9.**
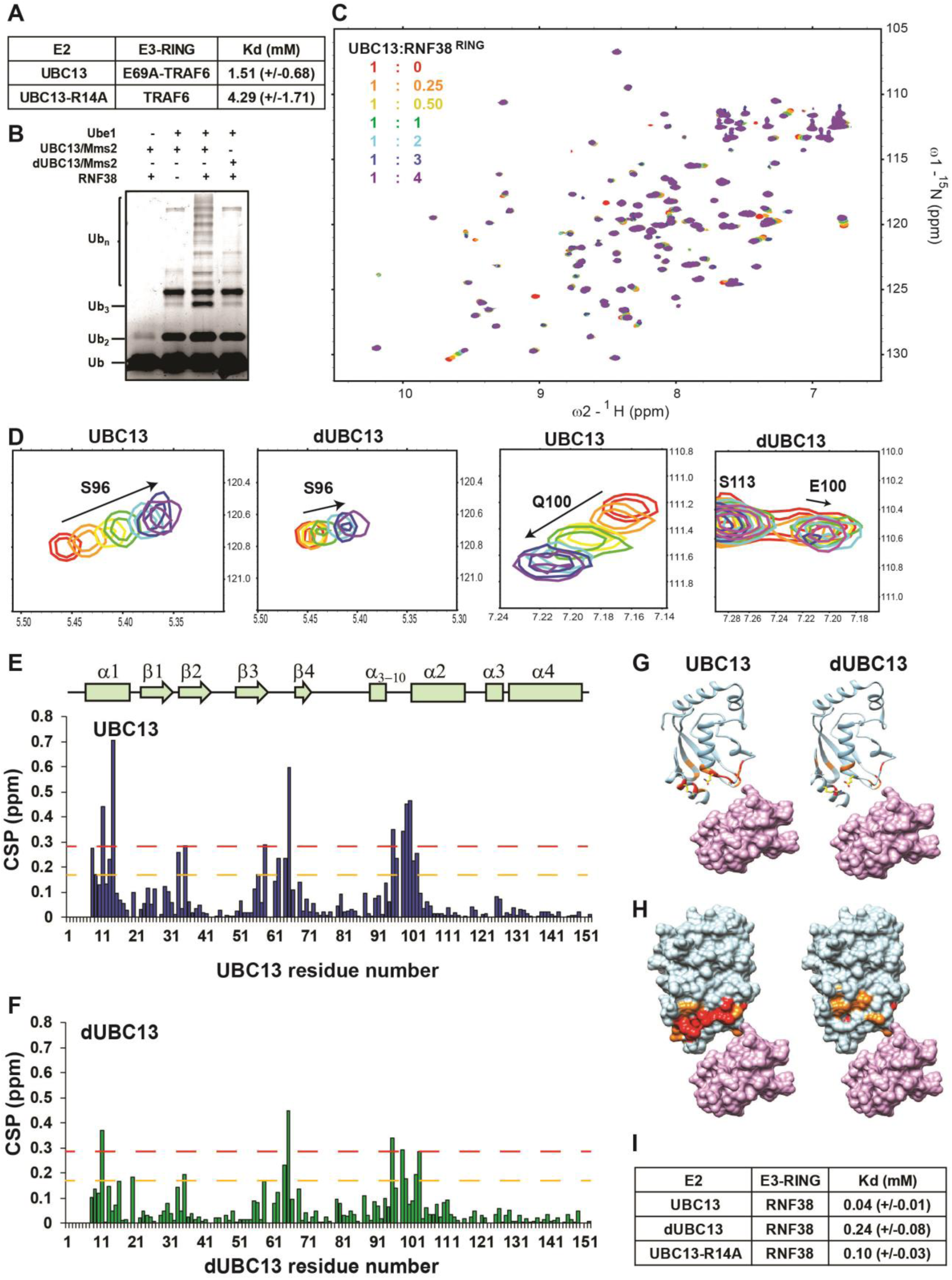
Activity and interactions of the UBC13/RNF38^RING^ complex. A) The measured dissociation constants of mutants of UBC13/TRAF6^RING^ complex are given as Mean+/−SD. B) *In vitro* ubiquitination assay was performed using Ube1, UBC13/Mms2 (or dUBC13/Mms2) and RNF38. C) Overlay of the ^15^N-edited HSQC spectra of free UBC13 (red) with different stoichiometric ratios of RNF38^RING^ as given in the top left-hand side of the spectra. D) Regions of the HSQC spectra are expanded to show the UBC13 and dUBC13 peaks during titration with RNF38^RING^. E) The CSPs for each residue in UBC13 upon binding to RNF38^RING^. The orange and red dashed lines correspond to Mean+SD and Mean+2*SD, respectively. The secondary structure alignment of UBC13 against its sequence is provided above the plot. F) The CSPs for each residue in dUBC13 upon binding to RNF38^RING^. The dashed lines are replicated from E). Significant CSPs were mapped on the UBC13 and dUBC13 structure using both the G) ribbon and H) surface representation. The UBC13 and dUBC13 are colored in light blue. The residues with CSPs above Mean+SD and Mean+2*SD are colored in orange and red, respectively. The RNF38^RING^ domain is surface rendered and colored in magenta. I) The measured dissociation constants of UBC13 and its mutants with RNF38^RING^ domain are given as Mean+/−SD.

### The effect of deamidation on transient interaction persists in the UBC13/RNF38^RING^ complex

Analysis of the UBC13/RING structures shows that R14-mediated salt-bridges are common in these complexes (Figure S13). An R14-mediated salt-bridge was present in four of the six complexes; CHIP, LNX1, TRIM25, and ZNRF1. However, in a few complexes like UBC13/RNF4 and UBC13/RNF8, the R14 mediated salt-bridge was absent. A typical RING domain from the E3 RNF38 (RNF38^RING^, aa: 387-465) was chosen to study the effect of UBC13 deamidation in such cases. RNF38^RING^ can activate UBC13 to synthesize polyubiquitin chains (Figure 9B). Similar to TRAF6, the RNF38^RING^ fails to activate dUBC13 (Figure 9B). In silico modeling of the UBC13/RNF38^RING^ native complex suggests that like RNF4 and RNF8, RNF38^RING^ lacks a negatively charged residue at the position corresponding to E69 in TRAF6^RING^ (replaced by L412, Figure S14). There are no negatively charged residues at the interface that can be an acceptor of the salt-bridge (Figure S14B). However, there are several acidic residues at the vicinity, similar to TRAF6^RING^, which can form non-native transient interactions with UBC13 (Figure S14C, S14D). In the absence of the intermolecular salt-bridge in the native complex, the loss of catalytic activity upon deamidation implies a drop in the UBC13/RNF38^RING^ affinity induced solely by perturbation of the transient complexes.

NMR titrations measured the effect of deamidation on the interaction between UBC13 and RNF38^RING^. When RNF38^RING^ domain was titrated into a sample of ^15^N-UBC13 (Figure 9C), major CSPs were observed in the α1-helix, β3-β4 loop, and the loop between α_3-10_ and α1-helix (Figures 9D and 9E). Unlike the UBC13/TRAF6^RING^ interface, the hydrophobic β3-β4 loop has a significant number of CSPs at the UBC13/RNF38^RING^ interface. Consequently, the K_d_ value of UBC13/RNF38^RING^ interaction was lower 0.04 (±0.01) mM (Figure 9I). Considerable peaks shifts were also detected when RNF38^RING^ was titrated to a sample of ^15^N-dUBC13 (Figure 9F). However, the CSPs in dUBC13 were reduced compared to UBC13 at the same protein: ligand stoichiometry, indicating reduced affinity (Figures 9F-H). Fitting of peak shifts against ligand: protein ratio yielded the K_d_ to be 0.25 (±0.08) mM, which corresponds to a 6-fold drop in affinity.

Both the effects of salt-bridge competition and repulsive interactions on transient complex could be at play in the dUBC13/RNF38^RING^ complex. The titration was repeated with R14A-UBC13, which mimics the salt-bridge competition in the transient complex. The K_d_ of R14A-UBC13/RNF38^RING^ interaction was 0.10 (±0.03) mM, corresponding to a 2.5-fold decrease in affinity exclusively due to the salt-bridge competition. Overall, the effect of deamidation on perturbing the ensemble of transient complexes could be verified in another RING domain, which indicated that the mechanism could be ubiquitous in UBC13/RING complexes.

### RING domains fail to bind and activate the dUBC13~Ub conjugate

If the RING domains bind weakly to dUBC13, their ability to activate the dUBC13~Ub conjugate should be compromised, which was tested by binding and activity assays. First, the conjugation of donor Ub to the UBC13 and dUBC13 were compared by an *in-vitro* conjugation reaction, where the E2s were incubated with E1, K63A-Ub, and ATP for 15 min. Both the UBC13 and dUBC13 conjugated with the donor Ub with similar efficiency, indicating that deamidation does not hamper the step of E2~Ub conjugation by E1 (Figure 10A and Figure S16). In the UBC13~Ub/TRAF6^RZ3^ complex, the RING domain and ZF1 domain makes additional contacts with donor Ub to stabilize the complex further. These contacts could rescue the loss in affinity between UBC13 and RING domain upon deamidation. Hence, the binding of UBC13~Ub and dUBC13~Ub to TRAF6^RZ3^ was compared. MBP-Mms2 was incubated with UBC13~Ub conjugate and TRAF6-RZ3 on amylose beads, washed thoroughly, separated on the gel and imaged. While UBC13~Ub conjugate bound to TRAF6^RZ3^, dUBC13~Ub conjugate failed to do so (Figure 10B and Figure S16). In a similar experiment, GST-RNF38^RING^ and UBC13~Ub were incubated on GST-beads, washed and separated on a gel. UBC13~Ub but not dUBC13~Ub bound to RNF38^RING^ (Figure 10C). Altogether, the binding experiments confirmed that secondary interactions between Ub/RING or Ub/ZF1 could not compensate for the loss of primary interaction between UBC13 and RING.

**Figure 10.**
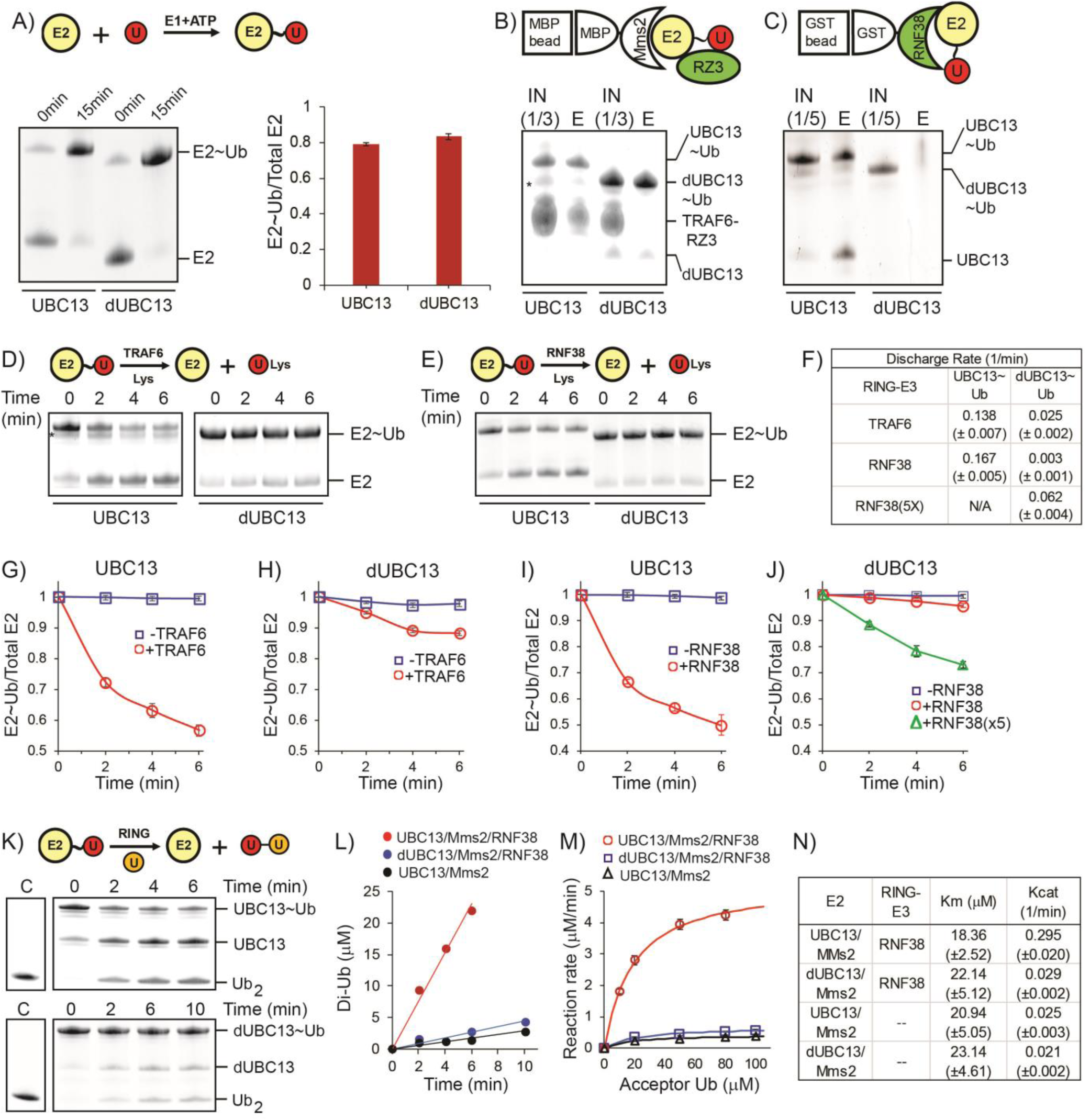
Activation of UBC13~Ub conjugates by RING domains. A) A comparison of Ub conjugation to UBC13 and dUBC13. E1, ATP, and UBC13/dUBC13 were incubated in reaction buffer for 15 min, quenched by adding EDTA and separated on SDS page. The amount of E2~Ub conjugates were quantified and plotted in the right section. The values are the mean of three reactions, and the error is the standard deviation of the same. B) Affinity pull-down experiment was performed by incubating MBP beads with MBP-Mms2, UBC13~Ub (or dUBC13~Ub) and TRAF6^RZ3^, washed thoroughly and separated on SDS gels. The asterisk denotes impurities. C) Affinity pull-down experiment was performed by incubating GST beads with GST-RNF38^RING^ and UBC13~Ub (or dUBC13~Ub), washed thoroughly and separated on SDS gels. D) Single-round discharge of Ub from UBC13~Ub and dUBC13~Ub catalyzed by TRAF6^RING^ was monitored. UBC13 was conjugated with Ub, and the reaction was quenched. Then Mms2, TRAF6^RING,^ and Lysine were added to the reaction mixture, and E2~Ub and free E2 was monitored over time. The proteins bands in D) were quantified and plotted in G) and H). E) Same as in D), where the discharge is catalyzed by RNF38^RING^ domain. The proteins in E) are quantified and plotted in I) and J). The rate of discharge in each case was calculated as discharge-rate= (Total E2-E2~Ub)/(Total E2.time) for the initial time points and given in F). K) The rate of Ub_2_ synthesis was monitored over time. UBC13 or dUBC13 was conjugated with Ub, and the reaction was quenched. Then Mms2, RNF38^RING^ and Acceptor Ub was added to the reaction mixture, and the synthesis of Ub_2_ was monitored over time. L) The rate of Ub_2_ synthesis for UBC13 and dUBC13 are compared at substrate (acceptor-Ub) concentration of 50 mM. M) The reaction rates for UBC13 and dUBC13 are compared at various substrate concentrations. N) The kinetic parameters of Ub_2_ synthesis for UBC13 and dUBC13 are given in a table. The Ub used in all the experiments of this figure is K63A-Ub, except the acceptor Ub used in K)-M) is D77-Ub.

The activation of UBC13~Ub conjugates by the RING domains were measured by the kinetics of single-round Ub discharge from the UBC13~Ub. UBC13 (or dUBC13) was first conjugated with K63A-Ub. Then the conjugation reaction was quenched, followed by the addition of Mms2, RING domains, and free Lysine. TRAF6^RING^ could effectively discharge Ub from UBC13~Ub, but not from dUBC13~Ub (Figure 10D, G, H). Similarly, RNF38^RING^ could discharge Ub from UBC13~Ub but not from dUBC13~Ub (Figure 10E, I, J). When the concentration of the RNF38^RING^ was increased by five-fold to compensate for the reduced affinity of dUBC13~Ub/RNF38^RING^ complex, the rate of Ub discharge improved by six-fold, which confirmed that the inefficient discharge is due to the reduced affinity of dUBC13~Ub/RNF38^RING^ complex (Figure 10K and 10F).

The lysine of the acceptor Ub at the growing end of a poly-Ub chain (either unanchored or anchored on a substrate), attacks the E2~Ub active site to transfer the donor Ub. The kinetics of single-round di-Ubiquitin (Ub_2_) synthesis was monitored to investigate if the deamidation of UBC13 affects the activity of UBC13 and its interaction with acceptor lysine at the active site. Here the lysine of the acceptor Ub mimics the acceptor lysine of growing poly-Ub chain. UBC13 (or dUBC13) was conjugated with donor K63A-Ub. Then Mms2, RNF38^RING^ and acceptor D77-Ub was added in the reaction mix, which was then monitored over time to detect the synthesis of Ub_2_. The rate of Ub_2_ synthesis was rapid for UBC13 but slow for dUBC13 (Figure 10L). Kinetic analysis indicated that the reaction rate for the dUBC13/RING complex is similar to the rate of UBC13 in the absence of RING (Figure 10M and Figure S17), indicating that the catalytic effect of RING is absent in the dUBC13/RING complex. A comparison of the kinetic parameters between UBC13 and dUBC13 indicated that deamidation does not significantly affect the Km of the substrate, but drastically affects the Kcat of the reaction (Figure 10N and Figure S17). This confirms that the area around the active site is not perturbed by deamidation, as was observed by NMR in Figure S2. Furthermore, in the absence of a RING domain, the Kcat of UBC13 and dUBC13 is similar, confirming that the catalytic activity of UBC13 is unaffected upon deamidation, similar to the observation of Figure 1C. However, unlike UBC13, the Kcat of dUBC13 does not increase in the presence of RING, confirming that RING domains fail to bind and activate the dUBC13~Ub conjugate.

## Discussion

Protein deamidation can have significant functional consequences for innate immune signaling (Zhao et al. 2016), which justifies its emergence as a powerful tool employed by pathogenic bacteria to suppress the immune response (Washington, Banfield & Dangl 2013). However, little is known about the molecular mechanism underlying deactivation of immune signaling by deamidation. The *Shigella flexineri* effector OspI suppresses the inflammatory response (NF-κB pathway) via deamidation of UBC13 at Q100. By an unknown mechanism, deamidation of Q100 to E100 inhibits the synthesis of K63-linked polyubiquitin chains by the UBC13/TRAF6 complex, preventing the downstream activation of NF-κB pathway. Q100 is present at the vicinity of UBC13/TRAF6 interface but does not form direct contact with TRAF6. We report that deamidation neither alters the fold nor the enzymatic activity of UBC13. NMR and kinetics studies indicate that deamidation does not alter the active site and the Km of acceptor lysine. Instead, it inhibits the interaction between UBC13 and RING domains to diminish the catalytic effect of the RING. The dUBC13/TRAF6 interaction is inhibited because deamidated E100 competes with an intermolecular R14/E69 salt-bridge at the UBC13/TRAF6^RING^ interface to form an intramolecular salt-bridge, which destabilizes the native complex. The new intramolecular salt-bridge also inhibits the long-range transient interactions formed by R14 with TRAF6^RING^. In addition, repulsive interactions between E100 and the negatively charged interface of TRAF6^RING^ destabilize the transient complexes. Cumulatively, these mechanisms reduced the affinity between UBC13 and TRAF6^RING^ by 17-fold. Consequently, its efficiency in activating the UBC13~Ub conjugate drops by 6-fold (Figure 10F). The effect of deamidation is also manifested in another UBC13/RING complex. Deamidation reduced the rate of Ub-discharge and the rate of Ub_2_ synthesis by several folds in the UBC13/RNF38^RING^ complex (Figure 10F and 10N). These results provide first insights into the molecular mechanism behind the inactivation of the Ubiquitin pathway by bacterial deamidation.

For several E3s like RNF38, the RING domain can function as a monomer (Buetow et al. 2015). For others like TRAF6, the RING domains function as dimers (Middleton et al. 2017a; Yin et al. 2009). Since deamidation perturbed the binding of UBC13 to both TRAF6 and RNF38, it is likely that deamidation inactivates UBC13 for both homo/hetero-dimeric RING-E3s. In dimeric RINGs like TRAF6, the proximal RING interacts with one unit of UBC13~Ub, and the distal RING interacts with the donor Ub of another UBC13~Ub (Figure S15). Additionally for TRAF6 dimer, the first zinc-finger following the RING domain of one protomer also interacts with the donor Ub of the UBC13~Ub bound to another protomer (Middleton et al. 2017b). However, the binding between UBC13 and TRAF6^RING^ is independent of dimerization (Yin et al. 2009), indicating that binding between UBC13 and proximal RING is primary for the rest of the contacts. Indeed, deamidation inhibited the binding of UBC13 to both TRAF6^RING^ and to the longer dimeric TRAF6^RZ3^, confirming that secondary interactions between Ub/RING or Ub/zinc-finger cannot compensate for the loss of primary E2/RING interaction.

Though the interface of a protein complex has numerous interacting residues, a few hotspot residues contribute significantly to the binding energy than the rest (Moreira, Fernandes & Ramos 2007). Hotspot prediction algorithms reported R14 to be a hotspot at the UBC13/TRAF6^RING^ interface (Meireles n.d.; Zhu & Mitchell 2011). Analysis of all the UBC13/RING (or U-box) structures shows that intermolecular salt-bridges involving R14 are present in several of the UBC13/RING complexes (Figure S13). Binding studies by isothermal titration calorimetry measured that R14A substitution reduced the binding energy of UBC13/ZNRF1^RING^ complex by ~1.2 kcal mol^−1^ (Behera et al. 2018). Similarly, our NMR titrations measured that R14A substitution reduced the binding energy by ~1.4 kcal mol^−1^ in the UBC13/TRAF6^RING^ complex. Altogether, R14 appears to be a hotspot at the UBC13/TRAF6^RING^ interface. Deamidation of UBC13 disrupts the salt-bridge between R14 and E69 due to competition from E100, which destabilizes the UBC13/TRAF6^RING^ complex. Recently, the salt-bridge competition or “theft” mechanism was observed in the binding switch of Raf Kinase Inhibitory Protein (RKIP) to either Raf-1 or GPCR-kinase 2 (Skinner et al. 2017). Phosphorylation of S153 in RKIP created a new salt-bridge between pS153 and K157, which disrupted the pre-existing salt-bridges involving K157 and resulted in the local unfolding of RKIP. The unfolding inhibited binding of RKIP to Raf-1 but promoted its binding to GPCR-kinase 2. Unlike RKIP, deamidation does not alter the local fold of UBC13. While intramolecular salt-bridges compete within RKIP, the competition is between an intramolecular and intermolecular salt-bridge in UBC13. As expected, the intramolecular salt-bridge is dominant and effectively outcompetes the intermolecular salt-bridge (Figure 5E).

Besides the hotspots, electrostatic substitutions at the vicinity of the interface can also increase protein-protein association (and affinity) by several folds (Selzer, Albeck & Schreiber 2000). Such substitutions would shift the equilibrium between native and transient associations (Volkov et al. 2010). The proper conformation of transient complexes is vital for two proteins to form a stable, productive complex (Pan et al. 2019; Schilder & Ubbink 2013). How PTMs affect native protein-protein interactions observed in the ground state structure of the complex is well appreciated. However, little is known about how PTMs can affect the higher energy transient protein-protein associations. PTMs like phosphorylation, eliminylation, and deamidation can change the surface electrostatics of a protein (Ribet & Cossart 2010) to modulate the conformation and population of transient complexes significantly. By a combination of all-atom simulations and experimental data, this study provides molecular details of how PTMs can modulate transient protein-protein associations to inhibit interaction. In the dUBC13/RNF38 complex, deamidation only alters transient associations. Nonetheless, this causes a 10-fold drop in the catalytic activity of the complex (Figure 10N), indicating that modifying the transient protein-protein associations can severely affect function.

The fate of the polyubiquitinated substrate depends on the linkage specificity of the conjugated polyubiquitin chain, which in turn depends on the specificity of the E2/E3 interaction. Rationally designed E2/E3 interactions can change the fate and function of cellular proteins (van Wijk et al. 2009). Hence, the interfacial contacts that determine E2/E3 specificities are a subject of intense research (Christensen, Brzovic & Klevit 2007; Soss et al. 2011; van Wijk et al. 2009, 2012). The strength and functionality of E2/E3 interactions have been typically tested by mutating the interfacial residues (Christensen, Brzovic & Klevit 2007; Das et al. 2009, 2013). This work provides evidence that transient interactions have an equally significant role in E2/E3 interaction and function. Rational design strategies will improve significantly by incorporating transient E2/E3 interactions.

In several cases, the native interactions cannot explain the functional implications of PTMs. This study introduces a new approach to understand the functional role of PTMs by considering their transient interactions. For example, spontaneous deamidation (non-enzymatic) is rampant in age-onset diseases, including neurodegenerative and ocular diseases. Deamidation of amylin accelerates its aggregation and amyloid formation (Dunkelberger et al. 2012). Crystallins deamidate with age, which promotes its aggregation to trigger cataract (Pande, Mokhor & Pande 2015). Crystallin surfaces are highly charged, and possibly deamidation alters their charge distribution to promote non-native associations leading to aggregation. However, the sites of deamidation on crystallins do not provide a clear mechanism of the process. Deamidation-induced modulation of transient protein-protein associations may explain the tendency of these proteins to aggregate.

The bacterial effector OspI from *S. flexineri* presents a fascinating case, wherein a pathogen employs subtle mechanisms like salt-bridge competition and modification of the transient protein-protein associations, to produce a remarkable cumulative effect of inhibiting protein-protein interaction, polyubiquitination, and the host immune response. Such a mechanism could be a ubiquitous mode of regulating cellular pathways by PTMs.

## Acknowledgments

The NMR spectra were collected at the NMR Facility of the National Centre for Biological Sciences. The authors thank Prof. Rachel Klevit and Prof. Danny Huang, Prof. Catherine Day for reagents, and Dr. Purushotham Reddy for help with data collection. This work was funded by the Tata Institute of Fundamental Research. R.D. acknowledges the DBT-Ramalingaswamy fellowship (BT/HRD/23/02/2006).

## Methods

### Initial structures and molecular modeling

The interfacial contacts of UBC13/TRAF6^RING^ were obtained from PDB entry 3HCU using the typical cut-offs in the contact analysis tool in UCSF Chimera (Pettersen et al. 2004). Initial structures of UBC13 and TRAF6^RING^ for MD simulations and association rate constant (k_on_) estimation were obtained from the PDB entry 3HCU (Chain A/B), which represents a model of the native complex. For MD simulations of TRAF6^RING^, a C-terminal truncated model comprising of residues 50-148 was used. Substitutions were introduced by replacing existing sidechain with the best aligning rotamer from the Dunbrack rotamer library (Dunbrack 2002) in UCSF Chimera (Pettersen et al. 2004). Electrostatic surface potentials of UBC13 and TRAF6^RING^ were calculated using the Adaptive Poisson-Boltzmann Solver (Baker et al. 2001). The in silico model of UBC13/RNF38^RING^ was generated by superposing RNF38^RING^ (PDB entry 4V3K, chain F) onto TRAF6^RING^ in the UBC13/ TRAF6^RING^ complex using the structural alignment tool in UCSF Chimera. The calculations based on transient-complex theory were performed using the TransComp webserver (Qin, Pang & Zhou 2011).

### General molecular dynamics (MD) simulation protocol

All simulation methodologies employed in the study were performed using the AMBER99SB-ILDN force field ((Best & Hummer 2009; Hornak et al. 2006; Lindorff-Larsen et al. 2010). Unbiased MD simulations were performed in GROMACS version 4.6.4, while biased simulations were performed using the pulled code in Gromacs 5.1.2 (Abraham et al. 2015; Hess et al. 2008). All the acidic and basic residues apart from Histidines were modeled in their charged states. The Zinc AMBER force field (Peters et al. 2010) parameters were used to model the two zinc coordination sites within the TRAF6^RING^ domain. The initial structures were solvated in an appropriate box using the TIP3P (Jorgensen et al. 1983) water model. The non-bonded ion parameters proposed by Joung and Cheatham (Joung & Cheatham 2008) were used to model Na^+^ and Cl^−^ ions in TIP3P water. Additional non-bonded parameter corrections proposed by Yoo and Aksementiev for cation-chloride (Yoo & Aksimentiev 2012), amine-carboxylate (Yoo and Aksimentiev, 2015) and aliphatic carbon-carbon (Yoo & Aksimentiev 2016b) interactions were used for all simulations to eliminate overestimation of the strength of these interactions. A suitable number of counterions were added to neutralize the residual charge of the system, and additional ions were added to the box depending on the desired concentration. The electrically neutral, solvated system was then subjected to energy minimization using the steepest descent method for a maximum of 5000 steps until the maximum force on any atom was less than 1000 kJ mol^−1^ nm^−1^. Production simulations were performed under periodic boundary conditions at a temperature of 300 K, and 1 bar pressure (NPT ensemble) following equilibration carried out for 600 ps with 2 fs time steps. Temperature control was achieved using the v-rescaling thermostat (Bussi, Donadio & Parrinello 2007) for both equilibration (*T_c_*= 0.1 ps) and production (*T_c_* = 2.5 ps) steps. The Berendsen (Berendsen et al. 1984) (*T_p_* = 1 ps) and Parrinello-Rahman (Parrinello & Rahman 1981) barostat (*T_p_* = 5 ps) were employed for pressure control during equilibration and production steps respectively. All bond lengths were constrained using the LINCS (Hess 2008) algorithm. Virtual interaction sites were employed for hydrogen atoms (Bjelkmar et al. 2010), which permitted the use of a 5 fs time step. Short-range electrostatics and van der Waals interactions were calculated using a 1.0 nm cut-off. Long-range electrostatics were calculated using the smooth Particle Mesh Ewald (PME) method (Darden, York & Pedersen 1993; Essmann et al. 1995). An analytical dispersion correction was applied to approximate the effect of long-range van der Waals interactions.

### Analysis of conventional MD simulations

Simulations of free dUBC13 and UBC13/dUBC13 native complexes with TRAF6^RING^ were performed using general MD protocol described above. Salt-bridge occupancies were computed using a cut-off distance of 0.5 nm between arginine Cζ and glutamate Cδ atoms using a python script. For unbiased-association simulations, UBC13/dUBC13 and TRAF6^RING^ were separated along with the x component of the vector connecting their center of masses by 4 nm in a cubic box with an edge length of 12.6 nm. This results in a box volume equal to ~2000 nm^3^ and the resulting concentration of each protein species equal to ~1 mM. Fifteen independent association runs (100 ns each with different initial atomic velocities) were performed at 10 mM NaCl and 100 mM NaCl concentrations.

Trajectories were analyzed every 250 ps using analysis scripts available within the Gromacs package. Minimum distances were measured between UBC13 and TRAF6^RING^ to identify native-like, non-native, and non-associating trajectories using g_mindist. Native-like transient complex formation was considered to occur in a trajectory when the minimum distance between the sidechain nitrogen atoms of R6/K10/R14 (UBC13) and sidechain oxygen atoms of D57/E69 (TRAF6 ^RING^) fell below 0.35 nm and persisted for more than 8 ns. Among the remaining trajectories, minimum distances were measured between all UBC13 and TRAF6 ^RING^ heavy atoms. Non-native and non-associating trajectories were then identified using the same distance and persistence period cut-off.

The spatial distribution of TRAF6^RING^ COMs around UBC13/dUBC13 was obtained by calculating the displacement between COMs of Cα atoms of helix-I in UBC13 and TRAF6^RING^ (70-109) using g_dist and plotted for all fifteen association trajectories. RMSD-based structural clustering (Daura et al. 1999) with a cut-off of 0.45 nm was performed on pairwise rmsd matrices (determined using g_rmsd) of the combined trajectories (1.5 μs) using UBC13/dUBC13 (Cα atoms) as a reference for superposition to obtain the UBC13/TRAF6^RING^ transient complex clusters. The rmsd was computed for the backbone atoms of both UBC13 and TRAF6^RING^ between all trajectory snapshots. The output structures of the complex were then superposed on UBC13 (PDB: 3HCU) to obtain the location of TRAF6^RING^ cluster representatives. Native complex formation was identified by measuring the Cα rmsd of TRAF6^RING^ with respect to its position in the crystal complex for trajectories which exhibited the formation of native-like transient complexes using UBC13 (Cα atoms) as a reference for superposition.

Two-dimensional free energy landscapes for UBC13/dUBC13 association with TRAF6^RING^ were computed as a function of minimum distance (for salt-bridges between UBC13-R6/K10/R14 and TRAF6-D57/E69) and TRAF6^RING^ rmsd using a python script. Bin width of 0.02 nm was used for both coordinates, and the free energy (*ΔG)* corresponding to each bin were determined using the relation:

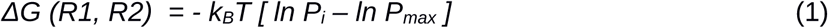

where R1/R2 are the two reaction coordinates, *k_B_*is the Boltzmann constant, T is the temperature, P_i_ is the joint probability of R1/R2 in a given bin and P_max_ is the max value of the probability. The lowest free energy state corresponds to ΔG = 0. Visualization of MD trajectories and analysis of the native contacts was performed using the MD analysis tool in UCSF Chimera.

### Umbrella sampling (US)

This is an enhanced-sampling method (Torrie & Valleau 1977) to determine free energy differences for state transformations by calculating the potential of mean force (PMF) as the function of a predefined reaction coordinate (*ξ*). This amounts to performing multiple, equilibrium MD simulations along with a range of *ξ* values and determining the corresponding free energy (*A(ξ)*). At points along *ξ* which are referred to as windows, a harmonic restraining potential (*w_i_)* is added to the original Hamiltonian in order to restrain the system close to a target value (*ξ_i_)*:

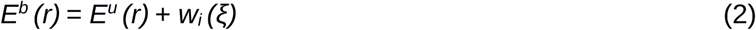

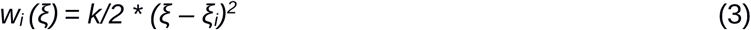

In equation (2), *E(r)* denotes the potential energy function (force field) of the system as a function of atomic coordinates (r). Subscripts b and u indicate biased and unbiased energies. *k is* the force constant for the harmonic bias potential whose magnitude determines the range of *ξ* values sampled within each window. The application of the bias potential particularly improves sampling in high energy regions along *ξ* which are poorly sampled in conventional MD simulations. The modified energy function yields biased probability distributions (P_i_^b^(ξ)) of *ξ* from which the unbiased, free energy (*A_i_(ξ))* of a window can be computed using the relation:

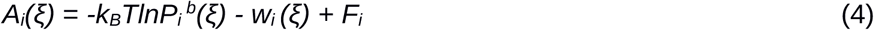

The value of F_i_ needs to be determined for all windows to combine adjacent windows and obtain the global free energy profile (A*(ξ))*, which describes state transformation. The weighted histogram analysis method (WHAM) (Kumar et al. 1992) is widely used to determine optimal values of *F_i_* in an iterative fashion so as to obtain convergence of the free energy profile.

Similar to previous studies which utilized the US approach to study the energetics of protein-protein association, the center of mass (COM) separation along the x-axis was chosen as the reaction coordinate to calculate the PMF for Barnase/Barstar and UBC13/TRAF6 ^RING^ association. Barstar and UBC13 were translated along the x-axis of the vector joining the COM between their binding partners to positions separated by 0.1 nm intervals to achieve final separations of 4.0 and 4.5 nm respectively. Each window was simulated in a rectangular box of dimensions 12.6 nm x 9.0 nm x 9.0 mm following equilibration. During production MD (10 ns for each window), a harmonic restraining potential (*k* = 1500 kJ mol^−1^ nm^−2^) was applied to the COM vector in each window and data were collected every 250 fs. The biased COM probability distributions from each window were analyzed based on WHAM and combined to obtain the global free energy (PMF) profile using the g_wham analysis tool (Hub, De Groot & Van Der Spoel 2010). A total of 200 bins were used to construct a mean PMF profile from five independent time blocks (1.5 ns each from 2.5 to 10 ns) for each window (Table S1). A tolerance of 10^−9^ was used to check for convergence. The free energy of association (ΔG_PMF_) along the COM coordinate was determined by setting the free energy profile to zero in the unbound state (4.0 nm for Barnase/Barstar and 4.5 nm for UBC13/TRAF6^RING^) and determining the value of the profile at mean COM separation in the native complex (2.3 nm for Barnase/Barstar and 2.7 nm for UBC13/TRAF6^RING^) window during production MD.

### Steered MD

This is a non-equilibrium, enhanced-sampling technique wherein a time-dependent biasing potential is applied on a selected atom/COM of atomic groups to induce perturbation in the system (Grubm ller, Heymann & Tavan 1996). This is achieved by means of a moving dummy atom attached to the pull group via a stiff harmonic spring. This method has been previously employed to study the unfolding of protein domains and the dissociation of protein-ligand complexes. In this study, we carried out force-induced unbinding of TRAF6^RING^ from UBC13 using the constant velocity pulling method along the x-axis to a COM separation of ~6 nm in 30 ns long simulations. The external force (*F_ext_*) acting on the pull group as the simulation progresses with time (*t*) is defined by the relation:

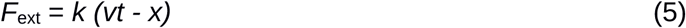

where *k* is the spring constant, v is the pull velocity, and x is the displacement of the ligand from its initial position. The pull force was applied to COM of Cα atoms of the RING domain (residues 70-109) at a low pull rate of 0.125 nm ns^−1^ (k=1500 kJ mol^−1^ nm^−2^). The same box dimensions were used as in the case of umbrella sampling. The COM of Cα atoms of UBC13 was kept fixed by periodically removing its rotational and translation motion to promote dissociation of the complex. The cumulative work (W) done to separate complex was calculated using the relation:

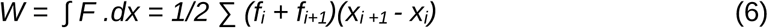

The work done over two successive simulation steps (i and i+1) is calculated as a product of the average force (f) multiplied by the displacement of the ligand between the steps. W is thus calculated by numerically integrating the work done between all successive steps of the simulation. The force and COM position of TRAF6^RING^ was recorded every 5 ps. F_max_ (rupture force) is defined as the maximum value of the force recorded in the Force-COM extension profile of an individual or averaged SMD trajectory. W was defined at a COM separation of 4.5 nm in the Work-COM extension profile as all native contacts were found to be disrupted at this COM separation. Both *F_max_*and *W* correlate with the binding affinity of the complex and hence allow for an analysis of the effect of mutations on the stability of the complex.

### Cloning and mutagenesis

A synthetic construct for TRAF6^RING^ (residues 50-124) was synthesized at Thermo Fischer Scientific, USA. PCR amplified ORF was cloned into a pGEX4T1 vector using NEB HF cloning kit (New England Biolabs, Inc. USA). E69A-TRAF6 was cloned using PCR based site-directed mutagenesis. Variants of UBC13 were generated using PCR based site-directed mutagenesis in WT-UBC13 in pET24 vector (a gift from Prof. Rachel Klevit). RNF38 plasmid was a gift from Prof. Danny Huang. TRAF6^RZ3^ plasmid was a gift from Prof. Catherine Day. All the plasmids were confirmed by sequencing.

### Protein expression and purification

All proteins were expressed in BL21 DE3 star cells (Invitrogen), grown at 37°C (in LB media for unlabeled proteins and in M9 media for C-13 or N-15 labeled proteins) till OD_600_ reached 0.7 and were induced with 0.5 mM IPTG for 4-5 hours. The harvested cells were lysed in 50 mM Tris, 250 mM NaCl, 5 mM βME, 2% glycerol, 0.01% triton-X, 1 mM PMSF and DNase at pH 7.5 using Emulsiflex high-pressure homogenizer. GST-TRAF6^RING^ and GST-RNF38^RING^ (residues 387-465) were bound to GSTrap HP columns (GE Healthcare) and were eluted with 10 mM reduced glutathione. The proteins were further purified by gel filtration in Superdex 75 pg 16/600 (GE Healthcare) in 25 mM Tris, 100 mM NaCl and 5 mM βME at pH 7.5. The His-tagged UBC13 and its variants were bound to HisTrap HP columns (GE Healthcare) and were eluted with 100mM-500mM Imidazole gradient. The proteins were further purified by gel filtration in Superdex 75 pg 16/600 (GE Healthcare) in 25 mM Tris, 100 mM NaCl and 5 mM βME at pH 7.5. TRAF6^RZ3^ was induced at OD_600_~0.7 with 0.2mM IPTG and 0.1mM ZnCl_2_. The lysed cells were resuspended in 50 mM Tris, 350 mM NaCl, 7mM βME, 1 mM PMSF, 10 mg Lysozyme, 0.01% Triton X, pH 8 and centrifuged. The supernatant was bound to HisTrap beads and was with 150mM-450mM Imidazole gradient. The eluent was subsequently purified by gel filtration in Superdex 75 pg 16/600 (GE Healthcare) in 25 mM Tris, 100 mM NaCl and 5 mM βME at pH 7.5

For the purification of Ub and its variants, lysed cells were resuspended in lysis buffer (50 mM Sodium acetate, 5 mM BME, 0.01% Triton X, pH 4.5), lysed by sonication and centrifuged to pellet cell debris. The supernatant was passed through the SP FF column (GE Healthcare) and the protein was eluted by gradient elution with increasing salt concentration (0 mM NaCl-600 mM NaCl). The proteins were further purified by gel filtration chromatography on Superdex 75 16/600 column (GE Healthcare) in 50 mM Sodium Phosphate, 100 mM NaCl, 5 mM BME, pH 6.5. For the purification of control K63-linked di-Ub, 1 µM UbE1, 40 µM Ubc13, 40 µM GB1-MMS2, 40 µM GST-RNF38, 200 µM D77-Ub and 200 µM K63A-Ub were mixed with reaction buffer (50 mM Sodium Phosphate, 5 mM MgCl_2_, 3 mM ATP, 0.5 mM DTT, pH 6.5) overnight. The mixture was dialyzed in 50mM Sodium Acetate, 5mM BME, pH 4.5, and centrifuged. The supernatant was passed through the SP FF column (GE Healthcare) and the di-Ub was separated by gradient elution with increasing salt concentration (0 mM NaCl-600 mM NaCl).

### NMR Spectroscopy

All NMR titration experiments were recorded at 298K on 600 MHz or 800 MHz Bruker Avance III HD spectrometer with a cryoprobe head. The samples were prepared in 25 mM Tris, 100 mM NaCl, pH 7.5, and 10% D_2_O. For NMR titration experiments, either ~2 mM TRAF6^RING^ or ~ 1 mM RNF38^RING^ were titrated into ~0.15 mM ^15^N-UBC13, ^15^N-dUBC13 and other mutants. The titration data was fit in 1:1 protein:ligand model using the equation CSP_obs_ = CSP_max_ {([P]_t_+[L]_t_+K_d_) - [([P]_t_+[L]_t_+K_d_)^2^-4[P]_t_[L]_t_]^1/2^}/2[P]_t_, where [P]_t_ and [L]_t_ are total concentrations of protein and ligand at any titration point.

The Arginine sidechain spectra of UBC13 and dUBC13 were recorded at 298K on 800 MHz Bruker Avance III HD spectrometer with a cryoprobe head, processed with NMRpipe (Delaglio et al. 1995) and analyzed with Sparky (Kneller & Kuntz 1993). The samples were prepared in 25 mM phosphate, 50 mM NaCl, pH 6.0. D_2_O was not directly added to the NMR sample to avoid additional signals due to 2H isotopomers. For NMR lock, the 3mm NMR tube was inserted co-axially in an external tube containing D_2_O. The arginine sidechain Hε, Nε, Cζ resonances were assigned by broadband NOESY-HSQC, HNCACB (Junji Iwahara and Marius Clore, JBNMR 2006), 2D HD(CD)NE and HD(CDNE)CZ (Frans Mulder JACS 2007) experiments.

### *In-vitro* ubiquitination assay, discharge assay, and Ub_2_ synthesis assay

For ubiquitination assay using TRAF6^RZ3^, E1 (0.5 μM), UBC13 and MMS2 (20 μM) and Alexa Fluor Maleimide (Invitrogen) labeled UbS20C (20 μM) were incubated with TRAF6^RZ3^ (15 μM) in UB-buffer (50 mM Tris, 5 mM ATP, 5 mM MgCl_2,_ 20mM NaCl, 2 mM DTT, pH 8) at 37°C for 10 minutes. The same reaction was repeated with GST-TRAF6^RING^ instead of TRAF6^RZ3^. The reaction mixtures were separated in 12% SDS gel, and the images were acquired in Uvitec (Cambridge). For ubiquitination assay using GST-RNF38^RING^, E1 (0.5 μM), UBC13 and MMS2 (5 μM) and Alexa Fluor Maleimide (Invitrogen) labeled UbS20C (20 μM) were incubated with GST-RNF38^RING^ (3 μM) in 20 mM Tris, 5 mM ATP, 5 mM MgCl_2_ (pH 7.5) at 37°C for 30 minutes. The reaction mixtures were separated in 12%-15% SDS gel, and the images were acquired in iBright FL1000 (Invitrogen). For Ub conjugation assays, E1 (1 μM), UBC13/dUBC13 (20 μM), and K63A-Ub (100 μM) were incubated in UB-buffer at 37°C for 15 minutes. For discharge assays, E1 (1 μM), UBC13/dUBC13 (20 μM), and K63A-Ub (100 μM) were incubated in UB-buffer at 37°C for 60 minutes. The reaction was quenched with EDTA (30 mM) at room temperature for 5 min. MBP-Mms2 (40 μM), TRAF6^RING^ or RNF38^RING^ (40 μM) and Lysine (15 mM) were added to start Ub-discharge. The reaction was quenched by 4X non-reducing SDS loading buffer at desired time-points. For Ub_2_ synthesis kinetics, E1 (1 μM), UBC13/dUBC13 (20 μM), and K63A-Ub (100 μM) were incubated in UB-buffer at 37°C for 60 minutes. The reaction was quenched with EDTA (30 mM) at room temperature for 5 min. Then MBP-Mms2 (40 μM), RNF38^RING^ (40 μM) and D77-Ub (20-100 μM) were added to start Ub_2_ synthesis. The reaction was quenched by 4X non-reducing SDS loading buffer at desired time-points. The reaction mixtures of Ub conjugation, Ub discharge, and Ub_2_ synthesis were separated in 12% SDS gel, stained overnight with SYPRO Ruby (Invitrogen) and the images were acquired in Uvitec (Cambridge).

### *In-vitro* binding assay

For the pull-down of TRAF6^RZ3^ with UBC13~Ub conjugate, an Ub conjugation reaction was performed as described above, and the reaction was quenched with EDTA at room temperature for 5 min. MBP-Mms2 (80 μM) was incubated on amylose bead (NEB) in 50 mM Tris, 20mM NaCl, pH 8. The UBC13~Ub conjugate (20 μM) was then mixed with TRAF6^RZ3^ (60 μM) and MBP-Mms2 on the amylose beads for 1 hour at 4°C, washed (3x, each time centrifuged at 3500 rpm for 3 min). After the final wash, the pellet was resuspended in 50 mM Tris, 20mM NaCl, pH 8 and separated on 12% SDS gel. For the pull-down of UBC13~Ub conjugate with GST-RNF38^RING^, the GST-RNF38^RING^ was incubated with glutathione agarose (Thermo Fisher). The UBC13~Ub conjugate was mixed with GST-RNF38^RING^ (60 μM) on glutathione agarose for 1 hour at 4°C, washed (3x), resuspended in 50 mM Tris, 20mM NaCl, pH 8 and separated on 12% SDS gel. The gels for both pull-down assays were stained overnight with SYPRO Ruby (Invitrogen), and the images were acquired in Uvitec (Cambridge).

