## Supporting Information for "Deamidation disrupts native and transient contacts to weaken the interaction between UBC13 and RING-finger E3 ligases"

**Table S1. $\Delta G_{PMF}$ values reported for the UBC13/TRAF6$^{RING}$ and the Barnase/Barstar complex.** Values are determined at 1.5 ns time intervals ranging from 2.5 to 10 ns. Related to Figures 3 and S6.

| $\Delta G_{PMF}$ (kcal mol$^{-1}$) | | | | | |
|---|---|---|---|---|---|
| **Time interval (ns)** | **UBC13** | **dUBC13** | **R14A - UBC13** | **Q100A - UBC13** | **Bs - Br** |
| 2.5 - 4.0 | -2.26 | -0.24 | 0.78 | -0.60 | -12.5 |
| 4.0 - 5.5 | -1.96 | -1.18 | -0.37 | -0.37 | -15.1 |
| 5.5 - 7.0 | -2.48 | 0.86 | 1.11 | -2.12 | -12.8 |
| 7.0 - 8.5 | -0.99 | -0.89 | -0.94 | -2.90 | -10.5 |
| 8.5 - 10.0 | -2.45 | -2.03 | -0.37 | -2.19 | -11.8 |

**Table S2. $F_{max}$ and unbinding work obtained from SMD trajectories of the UBC13/TRAF6$^{RING}$ and dUBC13/TRAF6$^{RING}$ complex**. Related to Figure 5 and Table 2.

| | **UBC13/TRAF6$^{RING}$** | | **dUBC13/TRAF6$^{RING}$** | |
|---|---|---|---|---|
| **Trajectory No.** | **$F_{max}$ (pN)** | **Work (kcal mol$^{-1}$)** | **$F_{max}$ (pN)** | **Work (kcal mol$^{-1}$)** |
| 1 | 666.9 | 38.6 | 659.2 | 27.3 |
| 2 | 698.1 | 39.5 | 612.4 | 34.6 |
| 3 | 615.7 | 44.9 | 658.8 | 25.7 |
| 4 | 585.1 | 41.3 | 513.6 | 24.7 |
| 5 | 714.1 | 43.1 | 503.3 | 22.4 |
| 6 | 642.9 | 30.4 | 518.6 | 34.0 |
| 7 | 644.5 | 32.0 | 683.6 | 38.0 |
| 8 | 628.3 | 28.4 | 592.7 | 28.9 |
| 9 | 558.0 | 32.1 | 574.6 | 27.9 |
| 10 | 808.7 | 49.8 | 529.9 | 32.4 |

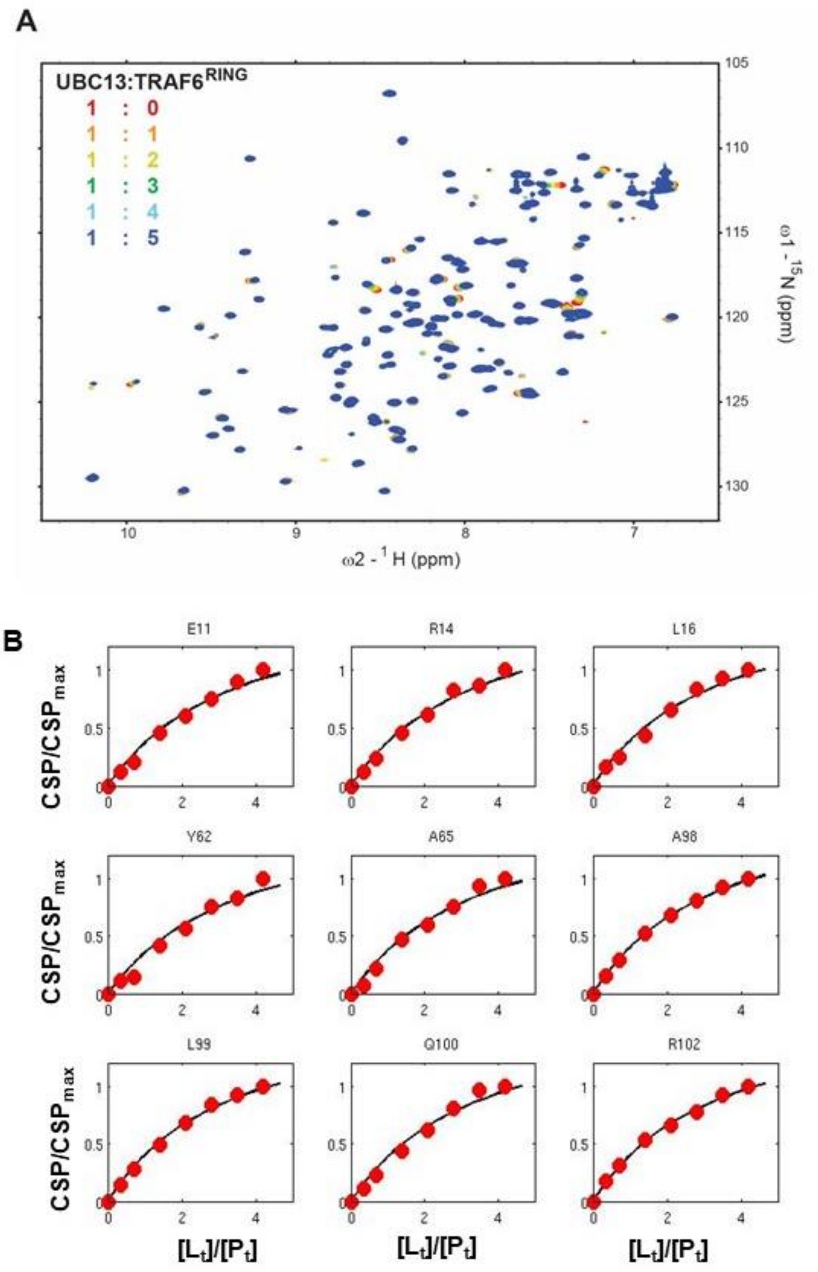


**Figure S1. NMR studies of UBC13 and dUBC13.**

Related to Figure 1.

A) A) Overlay of the $^{15}$N-edited HSQC spectra of free UBC13 (red) with different stoichiometric ratios of TRAF6$^{RING}$ as given in the top left-hand side of the spectra.

B) The fit of UBC13 peak shifts against the concentration ratio [TRAF6$^{RING}$]/[UBC13] ($[L_t]/[P_t]$) yielded the $K_d$ of the complex. The fit of nine typical residues is shown.

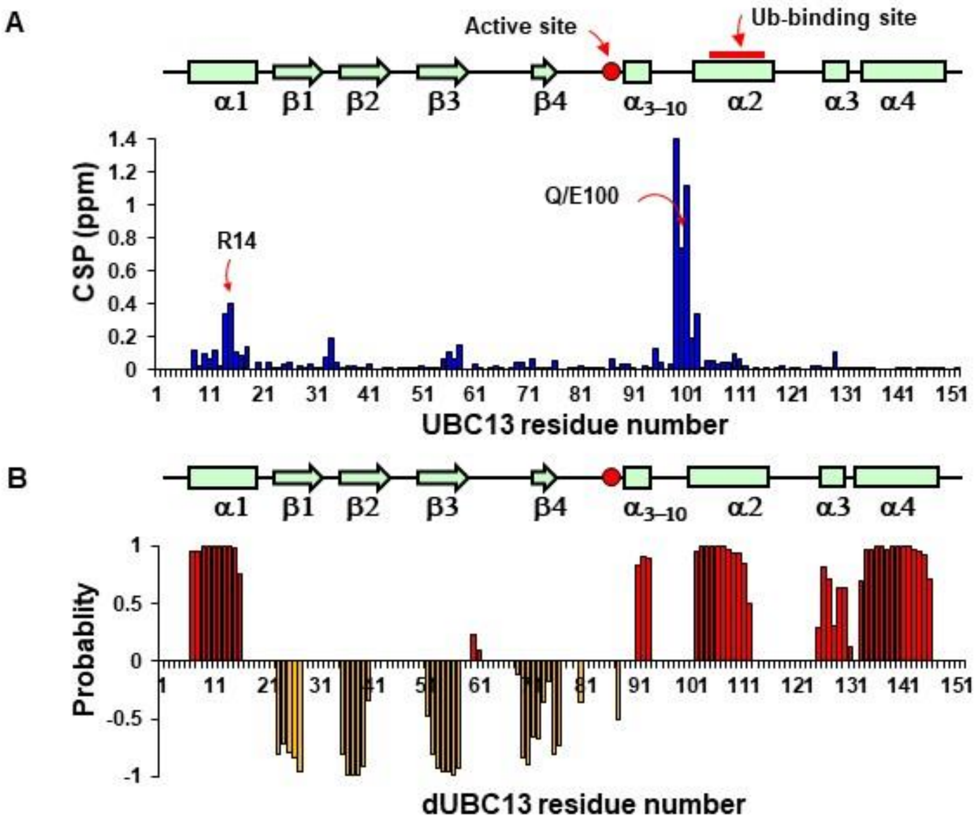


**Figure S2. NMR studies of UBC13 and dUBC13.**
Related to Figure 1.

A) The Chemical Shift Perturbations (CSPs) observed in amide resonances of UBC13 upon deamidation is plotted against the UBC13 residue numbers. The residues R14 and Q/E100 are shown by arrows. The linear arrangement of the secondary structural elements of UBC13 is indicated on the top for reference. The active site cysteine and the Ub-binding site are shown.

B) Secondary structure prediction of dUBC13 obtained by analyzing the backbone and $^{13}C_{\beta}$ chemical shifts by TALOS+(Shen et al. 2009). Positive values indicate α-helices, and negative values indicate β-strands. The linear arrangement of the secondary structural elements of UBC13 is shown on the top for reference.

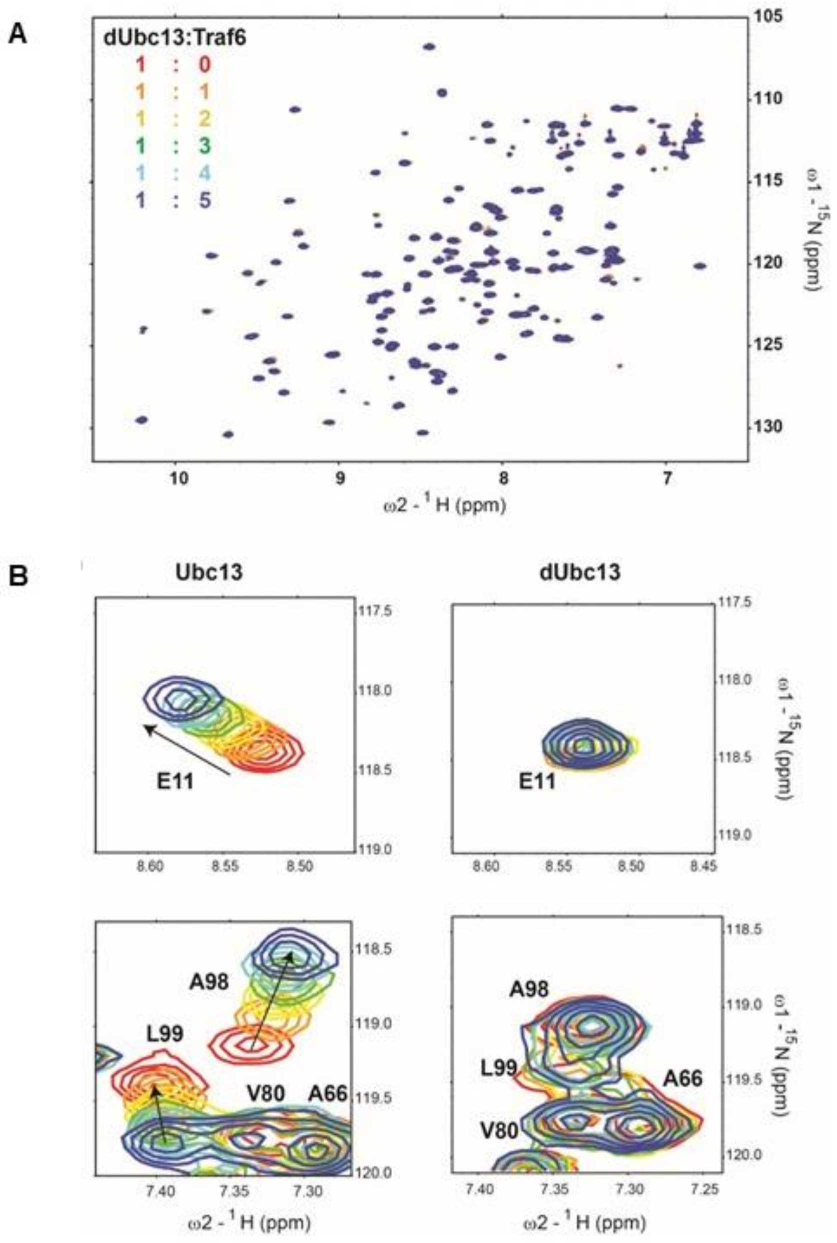


**Figure S3. Binding studies of dUBC13/TRAF6$^{RING}$ interaction.**

Related to Figure 1.

A) Overlay of the $^{15}$N-edited HSQC spectra of free dUBC13 (red) with different stoichiometric ratios of TRAF6 as given in the top left-hand side of the spectra.

B) Two regions of the spectra are expanded to show amide resonances (E11, A98, L99) that shift significantly in UBC13 but not dUBC13.

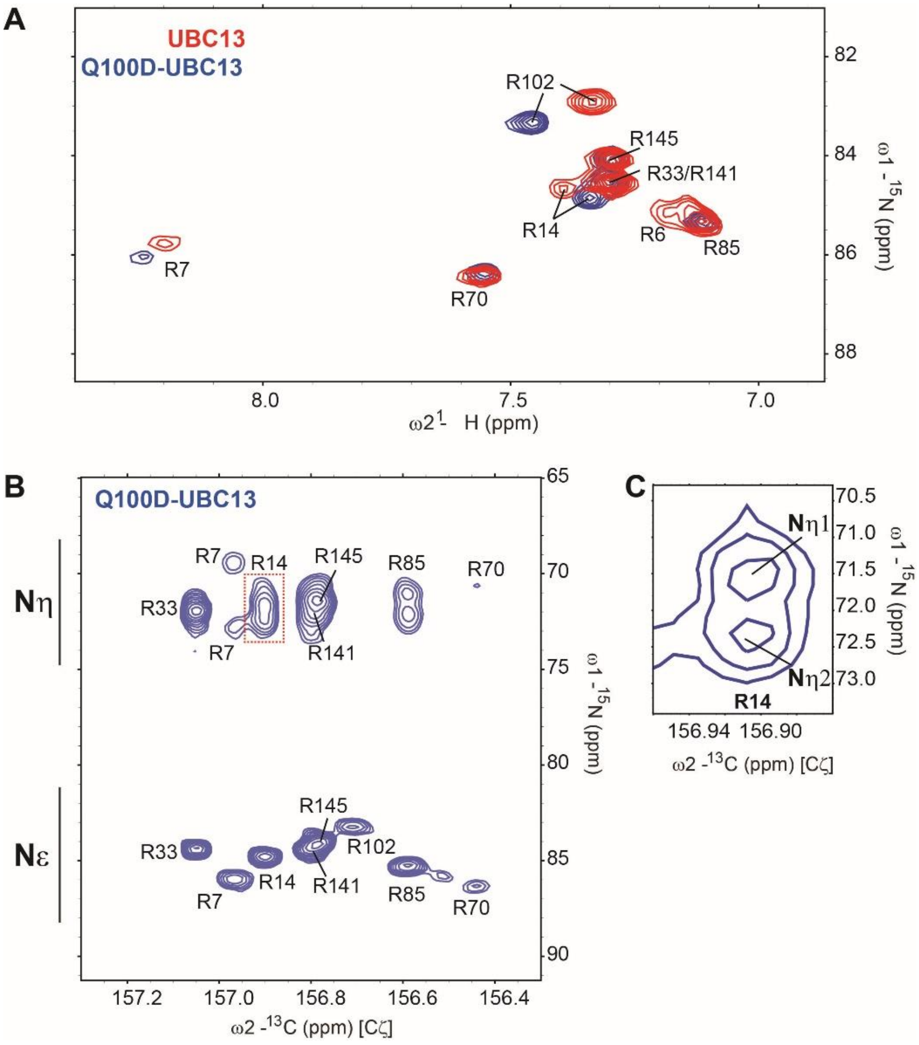


**Figure S4. Intermolecular salt-bridge persisted in Q100D-UBC13.**

Related to Figure 2 C-G

A) Overlay of UBC13 and Q100D-UBC13 $^{15}$N-$^{1}$H HSQC spectra zoomed around the Arginine Nε-Hε resonances, shows that R7, R14, and R102 sidechains resonances shift upon deamidation. B) The $^{15}$Nε/η-$^{13}$Cζ correlation spectra for Q100D-UBC13. The $^{15}$Nε/η and $^{13}$Cζ resonance shifts are in the x- and y-axis, respectively. C) $^{15}$Nε/η-$^{13}$Cζ correlation spectra for Q100D-UBC13 was processed with Gaussian window function and zoomed around the R14 $^{15}$Nη resonances to clearly show the splitting due to intramolecular salt-bridge.

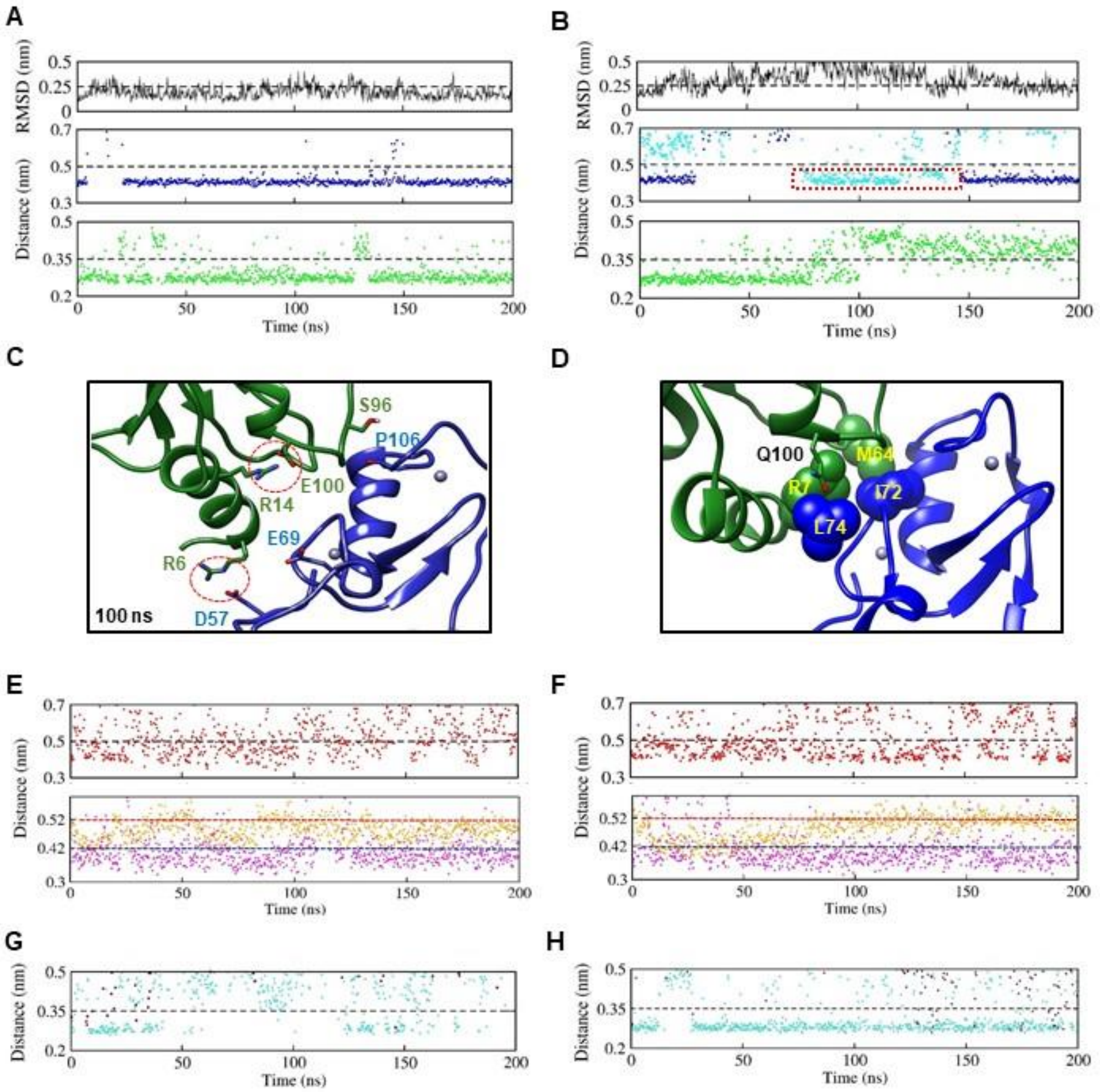


**Figure S5. Conventional MD simulations of the UBC13/ TRAF6$^{RING}$ and dUBC13/ TRAF6$^{RING}$ native complexes.**

Related to Figures 2B and 3.

A) RMSD and contact analysis of UBC13/TRAF6$^{RING}$ complex from a 200 ns MD simulation. B) RMSD and contact analysis of the dUBC13/TRAF6$^{RING}$ complex from a 200 ns MD simulation. For each trajectory in A and B, the top panel describes the RMSD variation of TRAF6$^{RING}$ with respect to the native complex as a function of time. The middle/bottom panels show the contact distance of two polar contacts against time: R14-Cζ/E69-Cδ (blue) and S96-Oγ/P106-N (green). The broken lines denote the corresponding distances observed in the

crystal structure plus 0.5 nm. In the middle panel of B, cyan dots depict the distance between the R14-Cζ and E100-Cδ atoms. The dotted box indicates the portion of the trajectory when the R14/E100 intramolecular salt-bridge formed.
C) A snapshot from the dUBC13/TRAF6$^{RING}$ native complex simulation at 100 ns showed that the R14/E100 salt-bridge could compete to disrupt the R14/E69 intermolecular salt-bridge.
D) The interactions between R7 and M64 from UBC13, and, I72 and L74 from TRAF6$^{RING}$ in the UBC13/TRAF6$^{RING}$ complex (PDB 3HCU) are shown.
E) The top panel shows the contact distance of R6/D57 salt-bridge (red) for the UBC13/TRAF6$^{RING}$ complex. The bottom panel describes the dynamics of two non-polar contacts R7/L74 (orange) and M64/L72 (magenta). The distances are calculated between R7-Cζ and I72-Cγ atoms for the R7/I72 interaction. The same is calculated between M64-Cε and L74/Cγ2 atoms for the M64/L74 interaction. The cut-offs are the distances observed in the crystal structure + 0.2 nm.
F) Same contacts as in E are analyzed for the dUBC13/TRAF6$^{RING}$ complex.
G) Variation in the contact distances of two hydrogen bonds K10/L74 (turquoise) and K94/N109 (maroon) against time are shown. Based on the significant variation in the contact distance in the simulations, these hydrogen bonds are not analyzed subsequently.
H) Same contacts as in G are shown for the dUBC13/TRAF6$^{RING}$ complex.

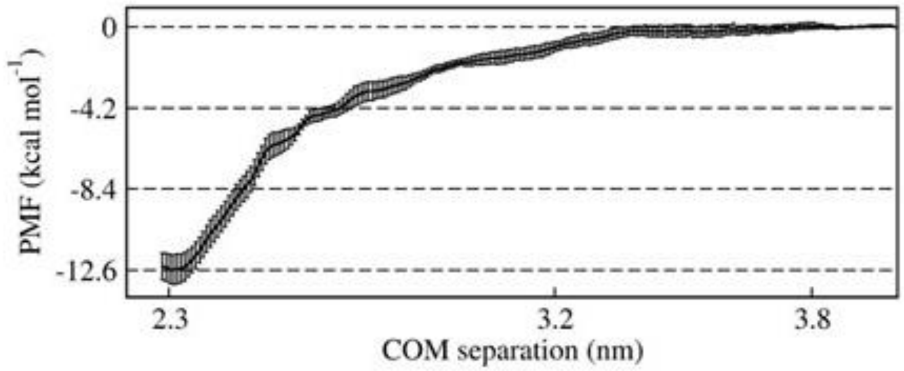


**Figure S6. Potential of mean force (PMF) profile for the Barnase-Barstar complex.**
The PMF of Barnase-Barstar complex is plotted against the Center of Mass separation. Related to Figure 3 and Table S1.

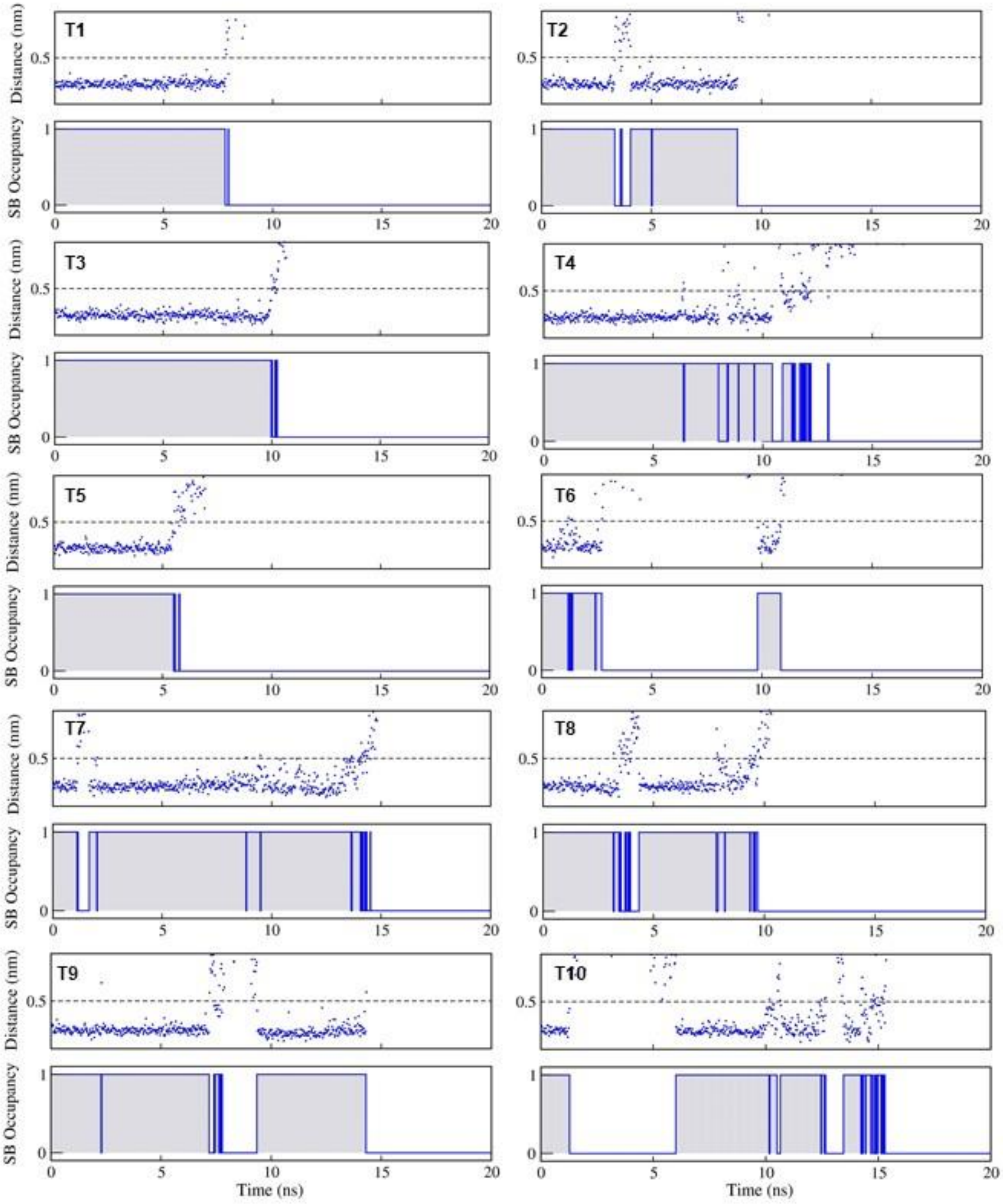


**Figure S7. R14-E69 salt-bridge dynamics across ten trajectories (T1-T10) during dissociation of the UBC13/TRAF6$^{RING}$ complex by SMD.**

Related to Figure 5 (A-C)**.**

For each trajectory, the actual distance variation and an occupancy plot are shown to indicate the presence (1) or absence (0) of the salt-bridge in a trajectory snapshot. The occupancies are obtained, as described in Figure 3A.

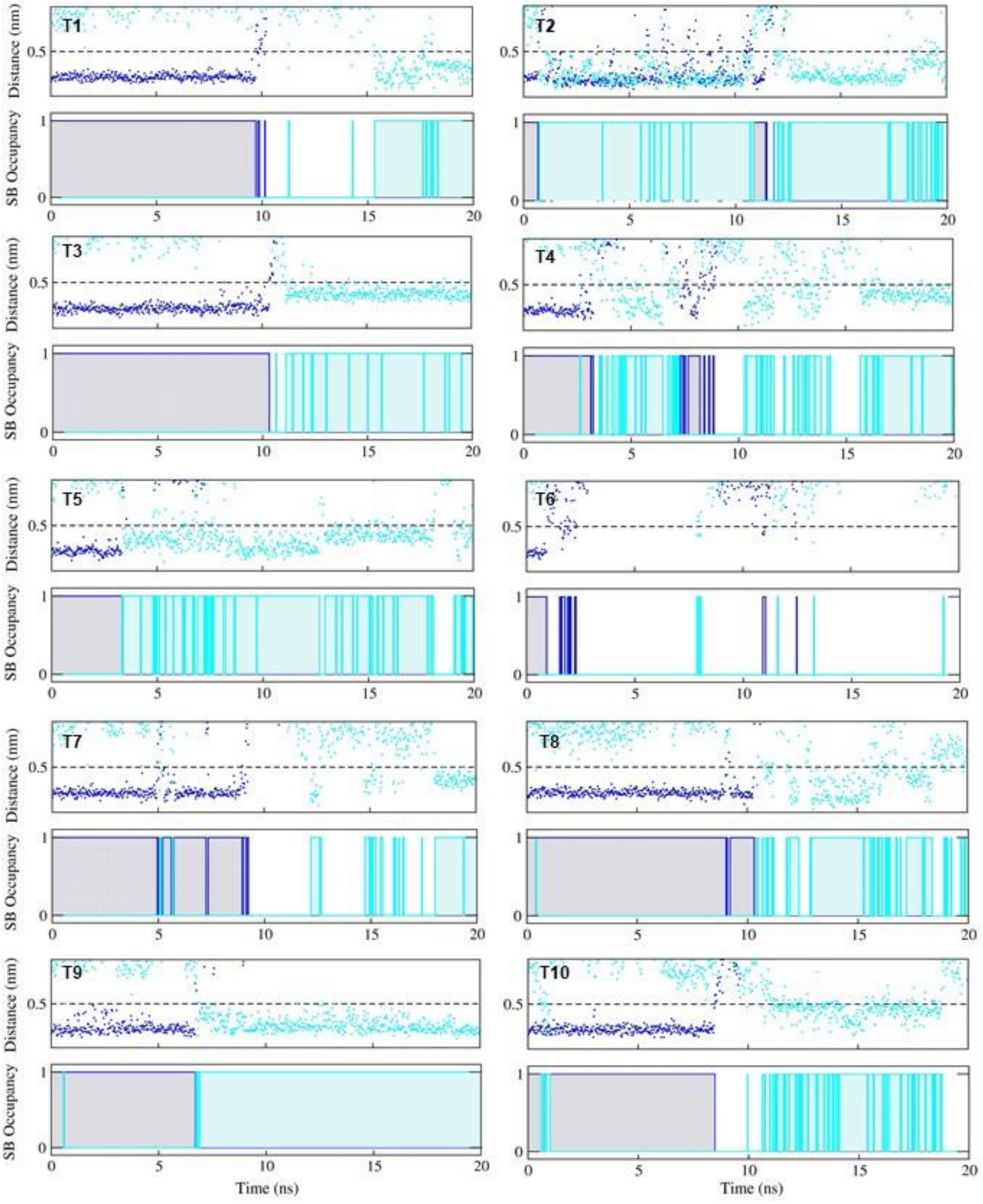


**Figure S8. R14-E69 (blue) / R14-E100 (cyan) salt-bridge dynamics across ten trajectories (T1-T10) during dissociation of the dUBC13/TRAF6$^{RING}$ complex by SMD**.
Related to Figure 5 (D-F).
For each trajectory, the actual distance variation and an occupancy plot are shown to indicate the presence (1) or absence (0) of the salt-bridge in a trajectory snapshot. The occupancies are obtained, as described in Figure 3A.

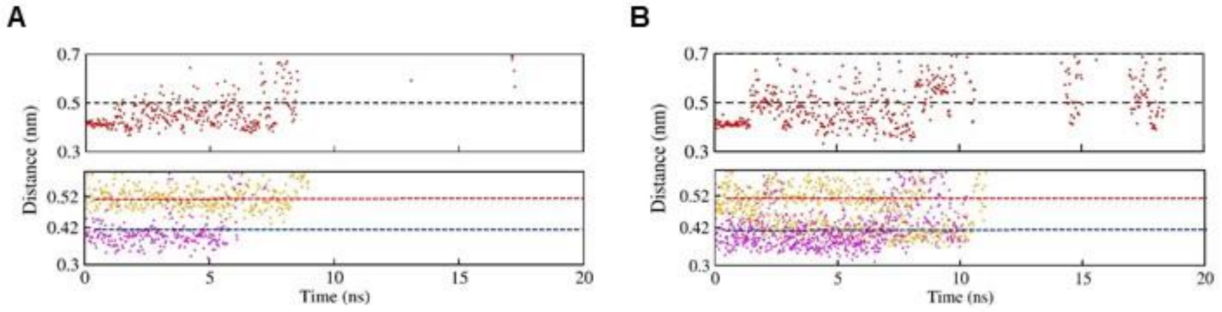


**Figure S9. Dynamics of R6-D57 salt-bridge and hydrophobic interactions in SMD.**
Related to Figure 5.
The contact distance of R6-D57 salt-bridge and hydrophobic contacts for A) UBC13/TRAF6$^{RING}$ and B) dUBC13/TRAF6$^{RING}$. In both A and B, the top panel describes the dynamics of R6-D57 salt-bridge (red, cut-off<=0.5 nm) while the bottom panel describes the dynamics of two non-polar contacts - R7-L74 (in orange, cut-off<=0.42 nm) and M64-L72 (in magenta, cut-off<=0.52 nm).

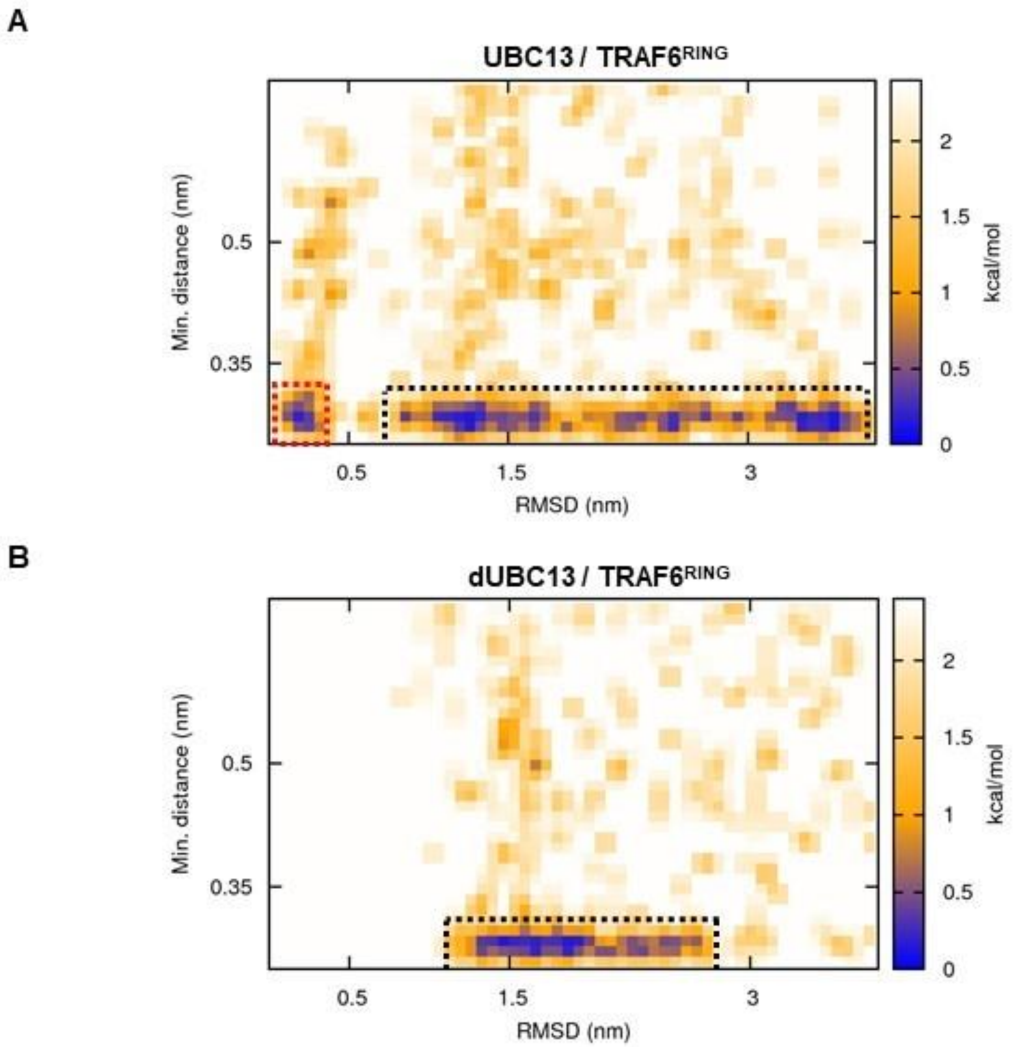


**Figure S10. Free energy landscapes obtained from association MD (10 mM NaCl).** Related to Figure 6 and 7**.**

A) and B) are 2D-free energy landscape of UBC13/TRAF6$^{RING}$ and dUBC13/TRAF6$^{RING}$ association, respectively. Both A and B are calculated as a function of minimum intermolecular salt-bridge distance (see Methods) and TRAF6$^{RING}$ RMSD with respect to its crystallographic orientation. Dotted black boxes indicate the native-like transient complex ensemble while the red dotted box in A indicates the native complex. Bins equal to or greater than 2.4 kcal mol$^{-1}$ are colored white.

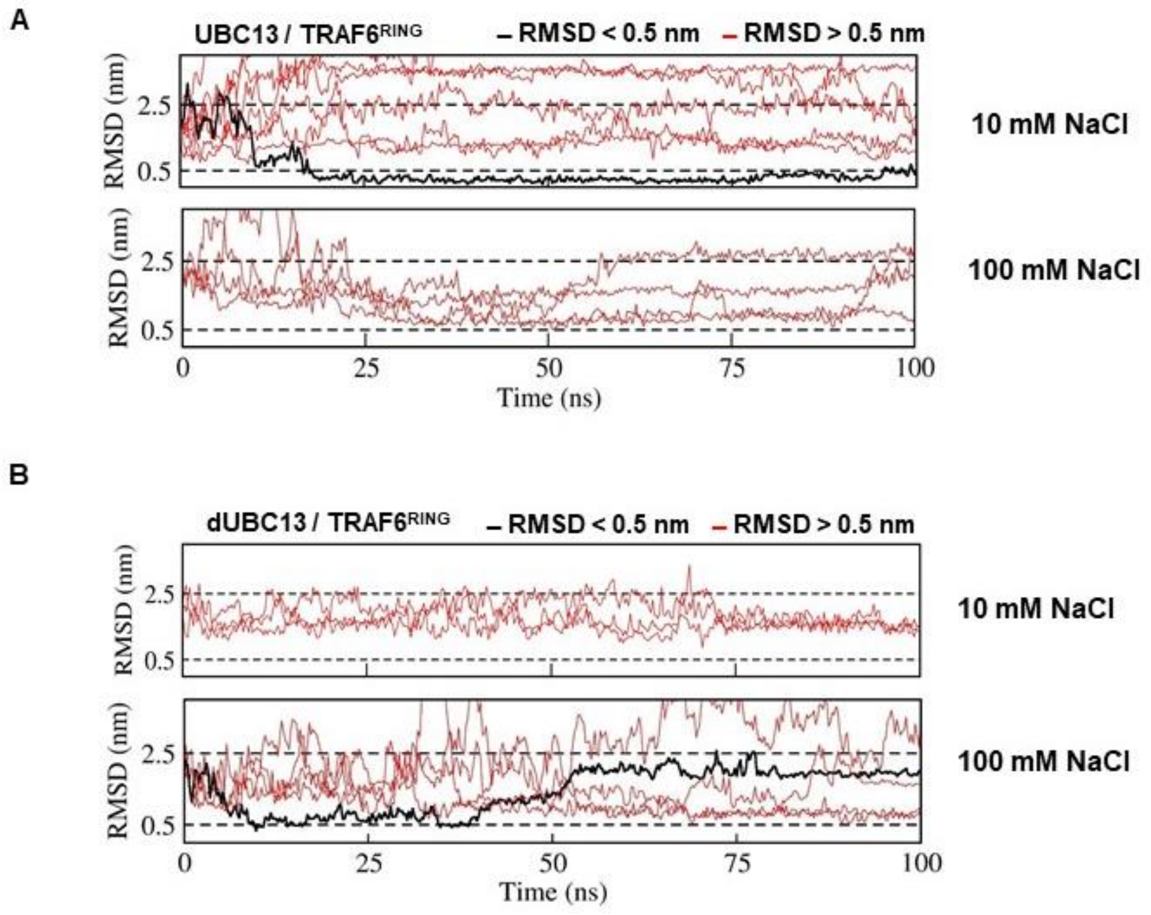


**Figure S11. RMSD analysis of UBC13/dUBC13 and TRAF6$^{RING}$ association trajectories.**

Related to Table 4 and Figures 6 and 7.

A) RMSD analysis of the UBC13/TRAF6$^{RING}$ trajectories, which show a native-like association. One trajectory at 10 mM NaCl showed TRAF6$^{RING}$ RMSD values close (~0.25 nm) to its crystallographic orientation. A detailed analysis of this trajectory is shown in Figure 7 (A, B) and Figure S12A. B) RMSD analysis of the dUBC13/TRAF6$^{RING}$ association trajectories, which show a native-like association. In only one trajectory at 100 mM NaCl (black lines), TRAF6$^{RING}$ RMSD values were transiently below 0.5 nm. A detailed analysis of this trajectory is shown in Figure 7 (C, D) and S12B.

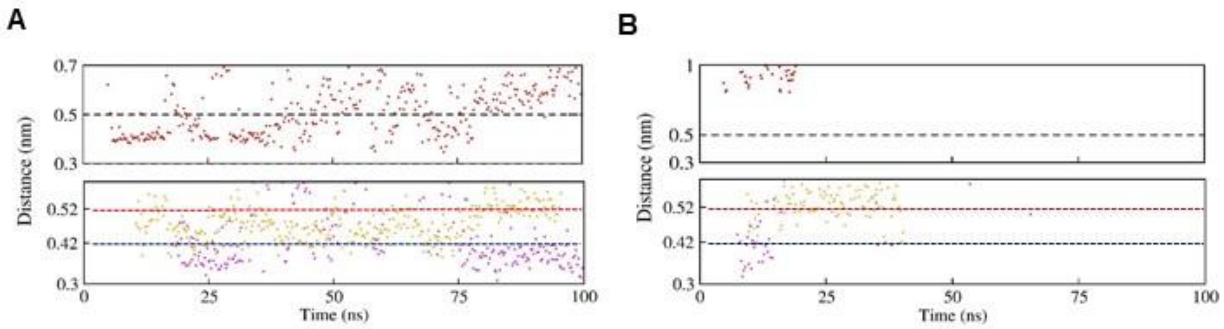


**Figure S12. Dynamics of R6-D57 salt-bridge and hydrophobic interactions in association MD.**

Related to Figure 8.

The variation in distance of R6-D57 salt-bridge and hydrophobic contacts for A) UBC13/TRAF6$^{RING}$ and B) dUBC13/TRAF6$^{RING}$. In both A and B, the top panel describes the dynamics of R6-D57 salt-bridge (red, cut-off<=0.5 nm) while the bottom panel describes the dynamics of two non-polar contacts - R7-L74 (in orange, cut-off<=0.42 nm) and M64-L72 (in magenta, cut-off<=0.52 nm). The distances measured are described in Figure S4E.

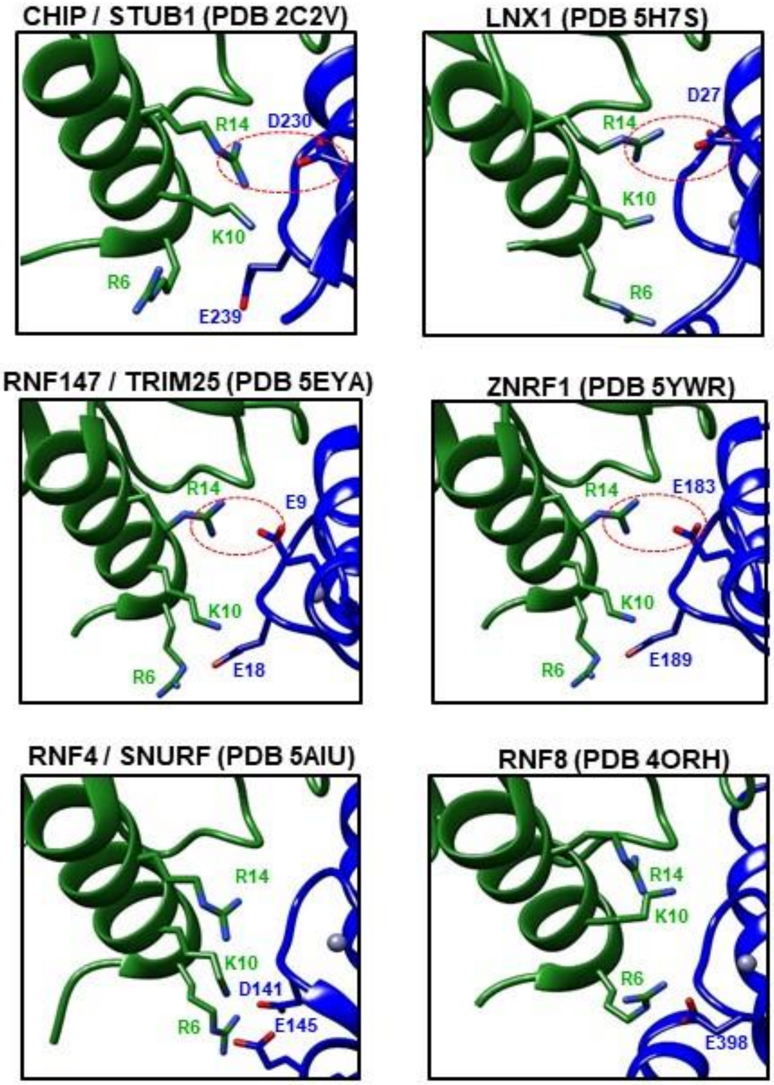


**Figure S13. R14-mediated intermolecular salt-bridges observed in crystallographic complexes of UBC13 with RING domains of other E3s.**

Dashed red circles are indicative of intermolecular salt-bridges involving R14 in UBC13. RNF4 and RNF8 lack R14-mediated intermolecular salt-bridges.

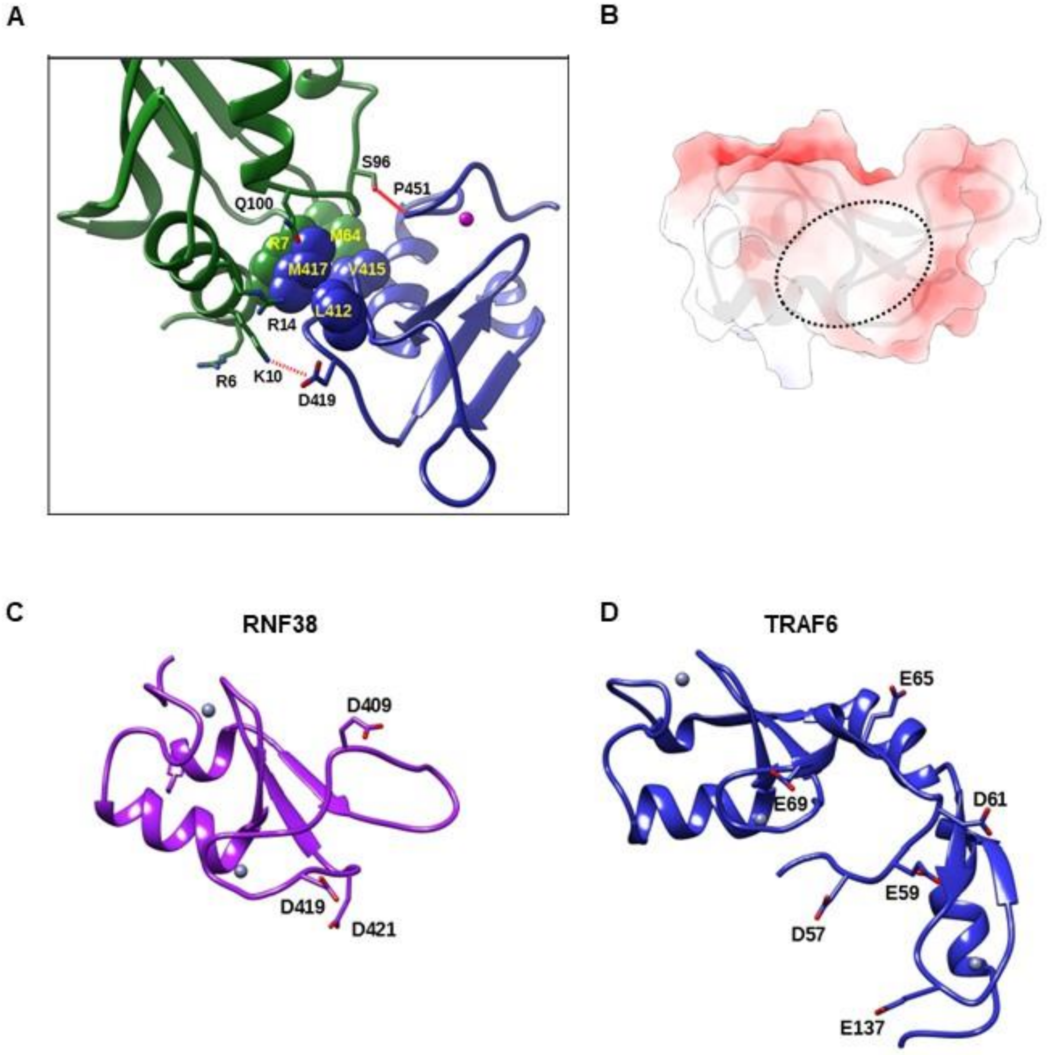


**Figure S14. Analysis of UBC13/RNF38$^{RING}$ *in silico* model and charge distribution at/near the association surface of RNF38/TRAF6$^{RING}$.**

Related to Figure 9.

A) *In silico* model of UBC13/RNF38$^{\mathbf{RING}}$. A weak polar bond between S96$^{UBC13}$ and P451$^{RNF38}$ is shown as a bold red line while a potential salt-bridge (no N-O hydrogen bonds observed) between K10$^{UBC13}$ and D419 $^{RNF38}$ is shown as a dashed red line. Residues participating in hydrophobic interactions are shown as spheres. The residue corresponding to E69$^{TRAF6}$ (which forms a salt-bridge with R14$^{UBC13}$) in RNF38 is L412. B) Surface electrostatic potential reveals a lack of salt-bridge acceptor at the RNF38$^{RING}$ interface in the complex shown in A). Colour scale varies from -8 to +8 $k_BT/e$. C) Negatively charged residues near the association surface of RNF38$^{RING}$. D) Negatively charged residues at/near the association surface of TRAF6$^{RING}$.

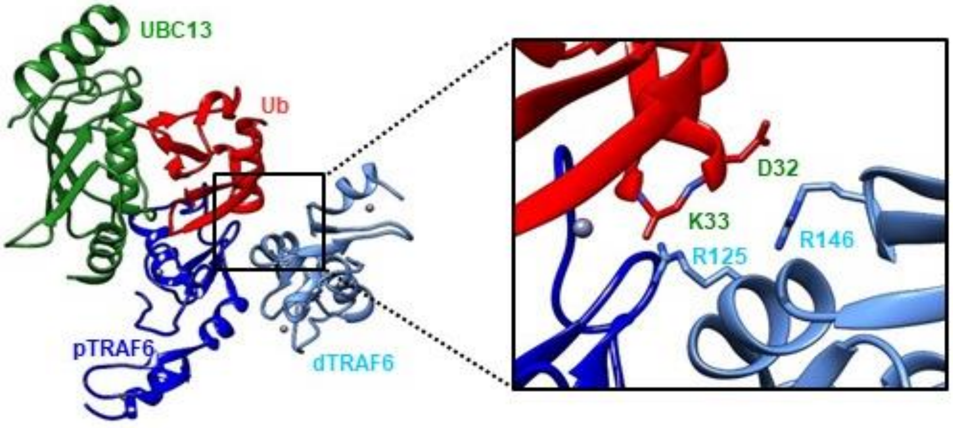


**Figure S15. The inability of the TRAF6$^{RING}$ homodimer to acquire the stereospecific orientation upon UBC13 deamidation prevents the stabilization of donor Ub.**

Structure of the UBC13~Ub-TRAF6$^{RING}$ catalytic complex (PDB 5VO0). The inset shows two essential Arginines located on the distal TRAF6 protomer (dTRAF6), which help to stabilize the closed conformation of UBC13~Ub in addition to the proximal unit (pTRAF6).

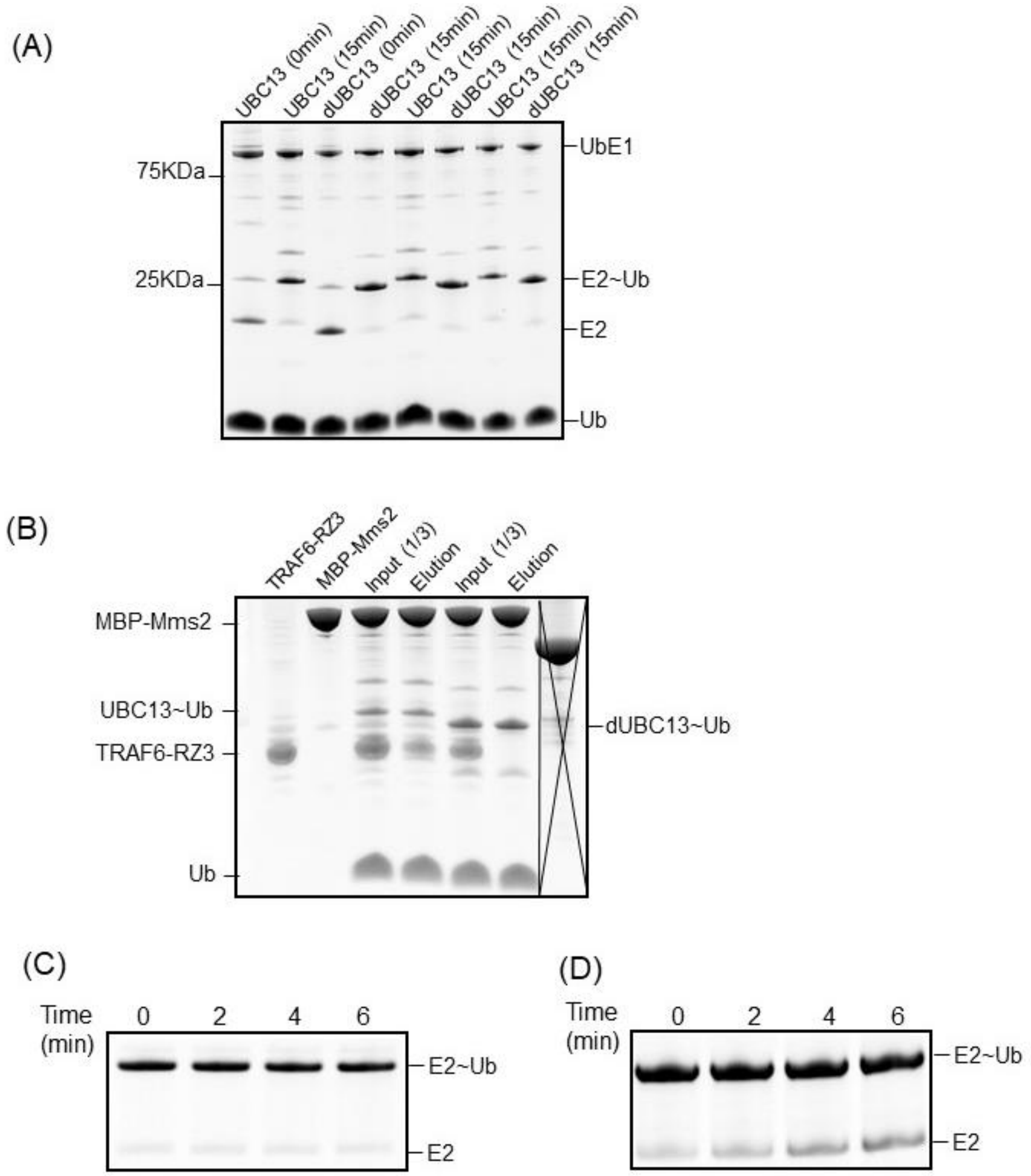


**Figure S16. Kinetics of Ub conjugation, RING-binding, and Ub-discharge.**

A) The complete gel-image of the Ub-conjugation reaction shown in Figure 10A. B) The complete gel-image of TRAF6-RZ3 and UBC13~Ub binding experiment of Figure 10B. C) The control UBC13~Ub discharge experiment in the absence of the RING domain. D) The dUBC13~Ub discharge experiment with RNF38(5X).

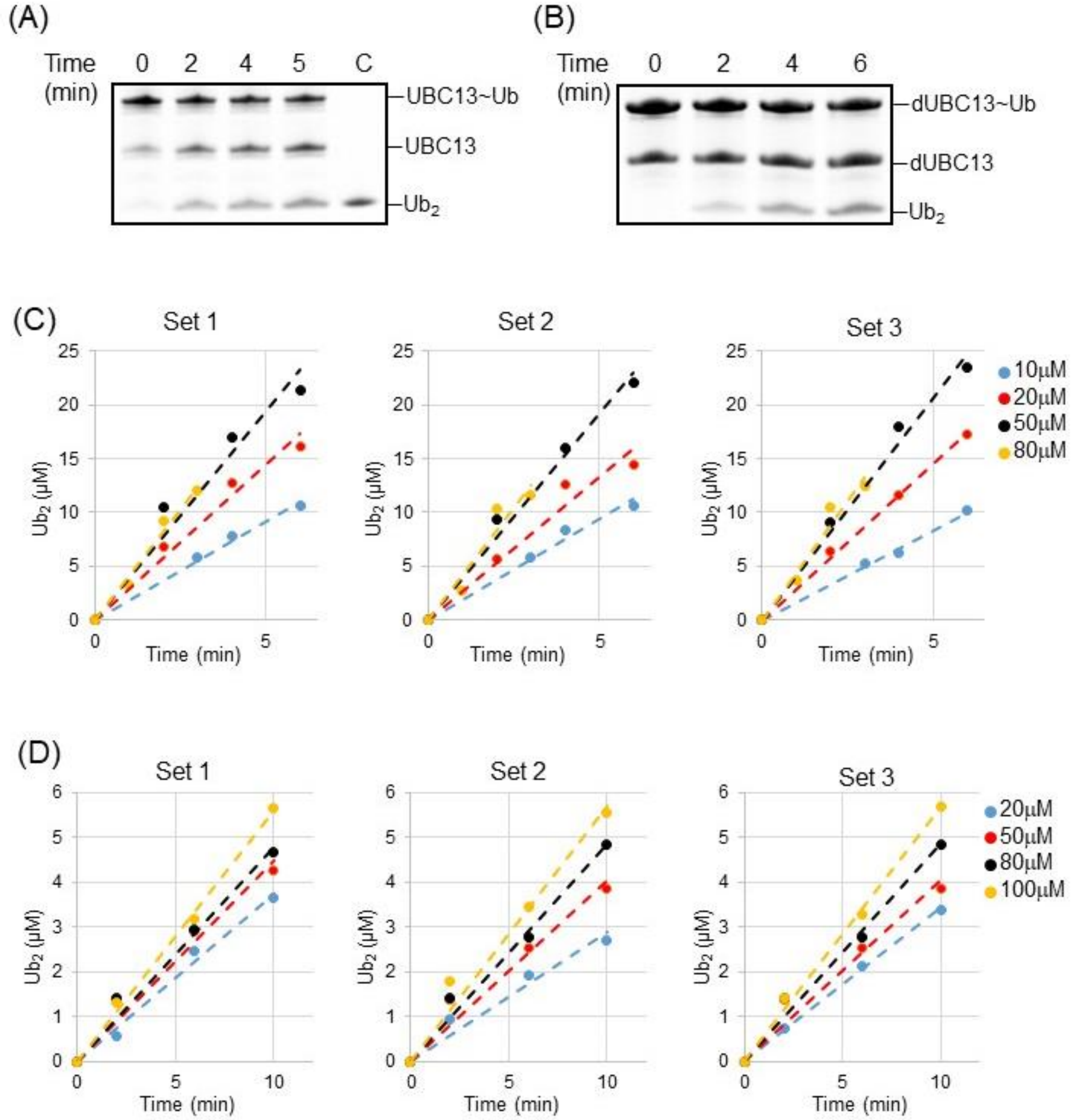


**Figure S17. Kinetics of $Ub_2$ synthesis.**

A) The kinetics of $Ub_2$ synthesis in UBC13/Mms2 the absence of RING domain. The control lane C is purified $Ub_2$. B) The kinetics of $Ub_2$ synthesis in dUBC13/Mms2 the absence of RING domain. The amount of $Ub_2$ formed by C) UBC13/Mms2/RNF38$^{RING}$ and d)dUBC13/Mms2/RNF38$^{RING}$ is plotted against time. The fits of the data are shown as dotted lines. The various substrate (D77-Ub) concentrations are shown by different colors.

**Supplemental Movies**

All movie files were prepared by pre-processing trajectories using a lowpass filter (g_filter tool in Gromacs) followed by the recording of trajectory snapshots in UCSF Chimera using the MD analysis module. In all trajectories, rotational and translational motion of UBC13 was removed entirely to fix its position with respect to $TRAF6^{RING}$.

**Movie S1 : Destabilization of the dUBC13/$TRAF6^{RING}$ native complex observed in a conventional MD simulation due to the formation of the R14-E100 intramolecular salt-bridge.** Related to Figure S5B.

The movie shows a disruption in the R14-E69 salt-bridge (~25 ns) followed by the formation of R14-E100 salt-bridge from ~75 to 150 ns (middle panel). This leads to a loss of the native intermolecular orientation as indicated by an increase in RMSD of $TRAF6^{RING}$ beyond 0.5 nm, as shown in Figure S5B (top panel). During this period, the S96-P106 contact is also lost (bottom panel).

Residues R6/R14/S96/E100 in UBC13 (Green) and D57/E69/P106 in $TRAF6^{RING}$ (Blue) are shown in stick representation. A black, dashed-line is drawn between R14 (Cζ) and E100 (Cδ) which indicates the presence/absence of an intramolecular salt-bridge depending on the length of the line.

**Movie S2 : Dissociation pathway of the UBC13/$TRAF6^{RING}$ native complex observed by steered MD.** Related to Figures 5 (A-C) and S9A.

The movie shows the order of contact disruption during enforced dissociation of $TRAF6^{RING}$ from UBC13. The loss of S96-P106 and hydrophobic interactions occur by ~9 ns. The R14-E69 salt-bridge persists until ~15 ns following after which, complete dissociation occurs.

Residues R6/R14/S96 in UBC13 (Green) and D57/E69/P106 in $TRAF6^{RING}$ (Blue) are shown in stick representation.

**Movie S3 : Dissociation pathway of the dUBC13/$TRAF6^{RING}$ native complex observed by steered MD.** Related to Figures 5 (D,E & F) and S9B.

The movie shows the order of contact disruption during enforced dissociation of $TRAF6^{RING}$ from dUBC13. The R14-E69 salt-bridge is lost within ~3 ns due to competition with E100 (Figure 5D, top panel). Loss of S96-P106 and hydrophobic interactions occur by ~11 ns. The R14-E100 intramolecular salt-bridge persists even after dissociation of the complex.

Residues R6/R14/S96/E100 in UBC13 (Green) and D57/E69/P106 in $TRAF6^{RING}$ (Blue) are shown in stick representation. A black, dashed-line is drawn between R14 (Cζ) and E100 (Cδ) which indicates the presence/absence of an intramolecular salt-bridge depending on the length of the line.

**Movie S4 : Association pathway of UBC13/TRAF6$^{RING}$ (10 mM NaCl) observed by conventional MD.** Related to Figures 8 (A,B) and S12A.

The movie shows the order of contact formation during the association of TRAF6$^{RING}$ and UBC13. Initial association occurs through the formation of R6-D57 and R14-E69 salt-bridge. Subsequent formation of hydrophobic and S96-P106 contacts lead to the acquisition of the native orientation (top panel, RMSD~0.3 nm) .

Residues R6/R14/S96 in UBC13 (Green) and D57/E69/P106 in TRAF6$^{RING}$ (Blue) are shown in stick representation.

**Movie S5 : Salt-bridge competition observed during dUBC13/TRAF6$^{RING}$ association (100 mM NaCl) by conventional MD.** Related to Figures 8 (C,D) and S12B.

The movie shows the order of contact formation during the association of TRAF6$^{RING}$ and dUBC13. Initial association occurs through the formation of a non-native R6-E69 salt-bridge due to the presence of the R14-E100 intramolecular salt-bridge. Hydrophobic and S96-P106 contacts are weakly present from ~5 to 40 ns. At ~22 ns, salt-bridge competition occurs, leading to the formation of the R14-E69 salt-bridge. E100$^{UBC13}$ eventually outcompetes E69$^{TRAF6}$ for R14 which leads to a disruption of the transient complex by 50 ns.

Residues R6/R14/S96/E100 in UBC13 (Green) and D57/E69/P106 in TRAF6$^{RING}$ (Blue) are shown in stick representation. A black, dashed-line is drawn between R14 (Cζ) and E100 (Cδ) which indicates the presence/absence of an intramolecular salt-bridge depending on the length of the line.
